# MSCs Successfully Deliver Oncolytic Virotherapy to Diffuse Intrinsic Pontine Glioma

**DOI:** 10.1101/2020.05.09.085837

**Authors:** Michael Chastkofsky, Katarzyna C. Pituch, Hiroaki Katagi, Liliana Ilut, Ting Xiao, Yu Han, Adam M. Sonabend, David T. Curiel, Erin R. Bonner, Javad Nazarian, Craig Horbinski, Charles D. James, Amanda M. Saratsis, Rintaro Hashizume, Maciej S. Lesniak, Irina V. Balyasnikova

## Abstract

Diffuse intrinsic pontine glioma (DIPG) is among the deadliest of pediatric brain tumors. Radiation therapy is the standard of care treatment for DIPG, but offers only transient relief of symptoms for DIPG patients without providing significant survival benefit. Oncolytic virotherapy (OV) is an anticancer treatment that has been investigated for treating various types of brain tumors. Here, we have explored the use of mesenchymal stem cells (MSC) for OV delivery and evaluated treatment efficacy using preclinical models of DIPG. Our results show that DIPG cells and tumors exhibit robust expression of cell surface proteins that are important for OV entry, and that MSCs loaded with OV disseminate within and release OV throughout the tumor in mice bearing DIPG brainstem xenografts. When combining administration of OV-loaded MSCs with radiotherapy, mice bearing brainstem DIPG xenografts experience a significant survival benefit, relative to that conferred by either therapy alone (p<0.0001). Our results support further preclinical investigation of cell-based OV therapy with radiation for potential translation in treating DIPG patients.

## Introduction

Diffuse intrinsic pontine glioma (DIPG) is an incurable pediatric brain tumor with a peak incidence between 6 and 9 years of age^1^. The current standard of care is irradiation, which temporarily relieves symptoms but has no significant effect on overall survival (OS), which is less than 18 months. Five-year survival for DIPG patients is less than 1%^2–5^. 80% of DIPG have mutation of genes encoding histone 3 (H3.1/3.3-K27M). Amplification of platelet-derived growth factor receptor alpha (*PDGFRA*), and mutations affecting activin A receptor type I (*ACVR1*) and tumor protein p53 (*TP53*), are also common among DIPG^6–9^ Due to their diffuse nature and location within the brainstem, DIPG are not surgically resectable^5^. As is the case for most brain tumors, systemically administered chemotherapeutics have limited access to DIPG due to the blood-brain barrier (BBB)^10–12^

The homing capacity of mesenchymal stem cells (MSCs) to tumors makes them excellent carriers of anticancer therapeutics. MSCs have been used as delivery vehicles for anticancer agents such as proteins, suicide gene/enzyme prodrugs, or oncolytic vectors, and are known for having tumor tropic properties^13–17^ MSCs can cross the BBB and reach brain tumor tissue following systemic administration. Intranasal delivery (IND) is an alternative route of therapy-carrying stem cells administration that has shown effectiveness in preclinical studies for treating neurological disorders including multiple sclerosis, viral brain infections, Parkinson’s disease, ischemic brain injury, and glioblastoma^18–22^.

We previously showed that the encapsulation of oncolytic virus (OV) CRAd.S.pK7 by MSCs has the potential to deliver a high therapeutic load to maximize tumor coverage with simultaneous protection from the fast clearance of the OV by the immune system^14^ Modification of the viral fiber in CRAd.S.pK7 with a heparin-binding domain, poly-L-lysine (pK7), increases the efficiency of OV entry into tumor cells^23^. Once in tumor cells, high level expression of viral proteins is driven by a survivin promoter^24^, which is active in numerous cancers, including GBM, but not in normal cells^25, 26^. Stem cell-mediated delivery of CRAd.S.pK7 ensures virus dissemination throughout the tumor. This OV delivery approach also delays elimination of the virus by the host immune system^14, 27^ and decreases neuroinflammatory response, thereby providing neuroprotection to normal brain^28^. The results from our preclinical studies in GBM models led to a phase I clinical trial investigating the safety of stem cell carrying OV for treating GBM patients (NCT03072134). We previously showed that oncolytic virotherapy, as administered locally to the tumor site or by IND using stem cell carriers, is an effective treatment when tested against models of glioblastoma. Stem cells are capable of delivering therapeutic cargos to distant parts of the brain^14, 22–24, 28–32^

In this study, we have evaluated CRAd.S.pK7 OV as a potential therapeutic for treating children with DIPG. CRAd.S.pK7 replication was examined in three patient-derived DIPG lines, and MSC migration as well as delivery of CRAd.S.pK7 was analyzed *in vivo* using DIPG brainstem xenograft models. Lastly, the efficacy of local and intranasal administration of MSCs carrying OV was determined through survival analysis of mice with brainstem xenografts, both in the presence and absence of radiation.

## Results

### In vitro and in vivo DIPG models

In order to investigate the potential of OV-carrying MSCs for DIPG treatment, we utilized a patient-derived xenograft (PDX) model of DIPG. Human DIPG cells were directly injected into the brainstem of mice (Fig. 1A) and formed tumors, as confirmed by three-dimensional bioluminescent imaging (BLI) and magnetic resonance imaging (MRI). Histologic analysis of resected mouse brains revealed an infiltrative growth pattern in the brainstem, consistent with that observed in patients (Fig. 1A).

**Figure 1.**
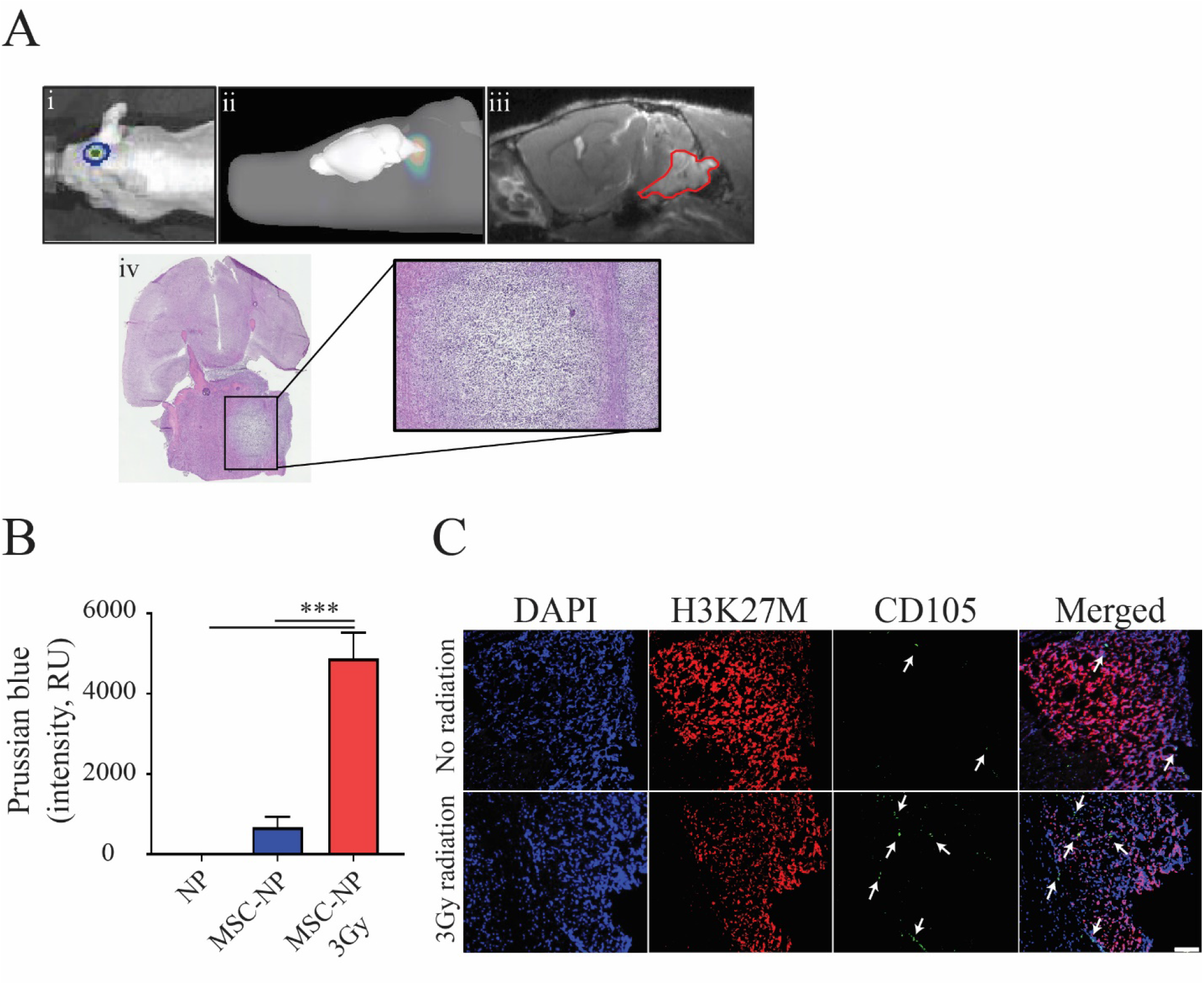
Radiation improves MSC migration toward DIPG tumors. A) Intracranial implantation of SF8628 DIPG cells results in pontine tumor formation in mice. Representative images show i) 2D-bioluminescent (BLI) image of the tumor, ii) 3D-rendered BLI, iii) MR image of tumor (outlined in red) depicting engraftment of the pontine tumor, and iv) histological verification of tumor by H&E stain. B) Irradiation significantly increased the trafficking of iron NP-labeled MSCs to the tumor site (One-way ANOVA with Tukey’s post-hoc test, n≤3. Level of significance: *** p <0.001). C) MSC migration to DIPG tumors was further validated using IHC (nuclear staining DAPI (blue), histone H3K27M (red) and MSCs-CD105 (green, indicated by white arrows). Scale= 100 μm.

### In vitro and in vivo effects of radiation on MSC tropism for DIPG

Initially, we investigated the effects of radiation on DIPG cells in culture and treated human SF8628 DIPG cells with fractionated irradiation for three days, resulting in the cumulative dose administration of 3, 6, and 12 Gy (SFig. 1A). Treatment of cells with 6 and 12 Gy irradiation revealed substantial reductions in viability in at days 5 (at 6Gy: 90.2±5.6%; at 12Gy:73.1±0.54%) and 6 (at 6Gy:19.26±6.7%; at 12Gy:29.38±6.7 6%))(SFig. 1A).

DIPG cells were treated with the sublethal dose of 3 Gy to examine the effect of radiation on MSC tropism to DIPG *in vitro*. MSC migration toward irradiated DIPG cells was significantly increased in relation to non-irradiated control cells (142.3±1.2% vs. 100±4.7%) (SFig 1B). Next, MSCs labeled with iron nanoparticles (SPIO) were administered by IND to mice with brainstem DIPG xenografts to evaluate radiation-associated effects on MSC migration to tumor *in vivo*. Prussian Blue staining revealed a significant increase in iron-labeled MSCs within and around brainstem tumor in irradiated animals, as compared to non-irradiated mice treated with iron nanoparticles alone (Fig. 1B, One-way ANOVA, p<0.0001). The tumor location of MSCs was confirmed by staining of tissue with anti-CD105 antibody (Fig. 1C). Collectively, these data demonstrate that fractionated low-dose radiation increases the migration of MSCs toward DIPG tumors.

### Irradiation increases the expression of chemoattractant cytokines in DIPG tissue

Tumor tissue was dissected from irradiated and non-irradiated mice to investigate potential mediators of MSC tropism for DIPG (SFig. 1C). Results from array analysis revealed multiple chemokines/cytokines as being upregulated in DIPG tumors in response to radiation treatment (Fig. 2A). Binding partners for two of these, bone morphogenetic protein 4 (BMP-4) and betacellulin (BTC), are expressed by MSCs, BMP receptor (BMPR) 2 and epidermal growth factor (EGFR) receptor, respectively (Fig. 2B-C). The siRNA suppression of BMP-4 or BTC alone in SF8628 cells (Fig. 2D) did not affect MSC migration toward conditioned media of irradiated DIPG cells. In contrast, concomitant silencing of both chemokines significantly reduced MSC tropism for radiation-treated DIPG cells (Fig. 2E-F, SFig. 1D). These data indicate that combined BMP4 and BTC expression is important for MSC tropism to irradiated DIPG tumors.

**Figure 2.**
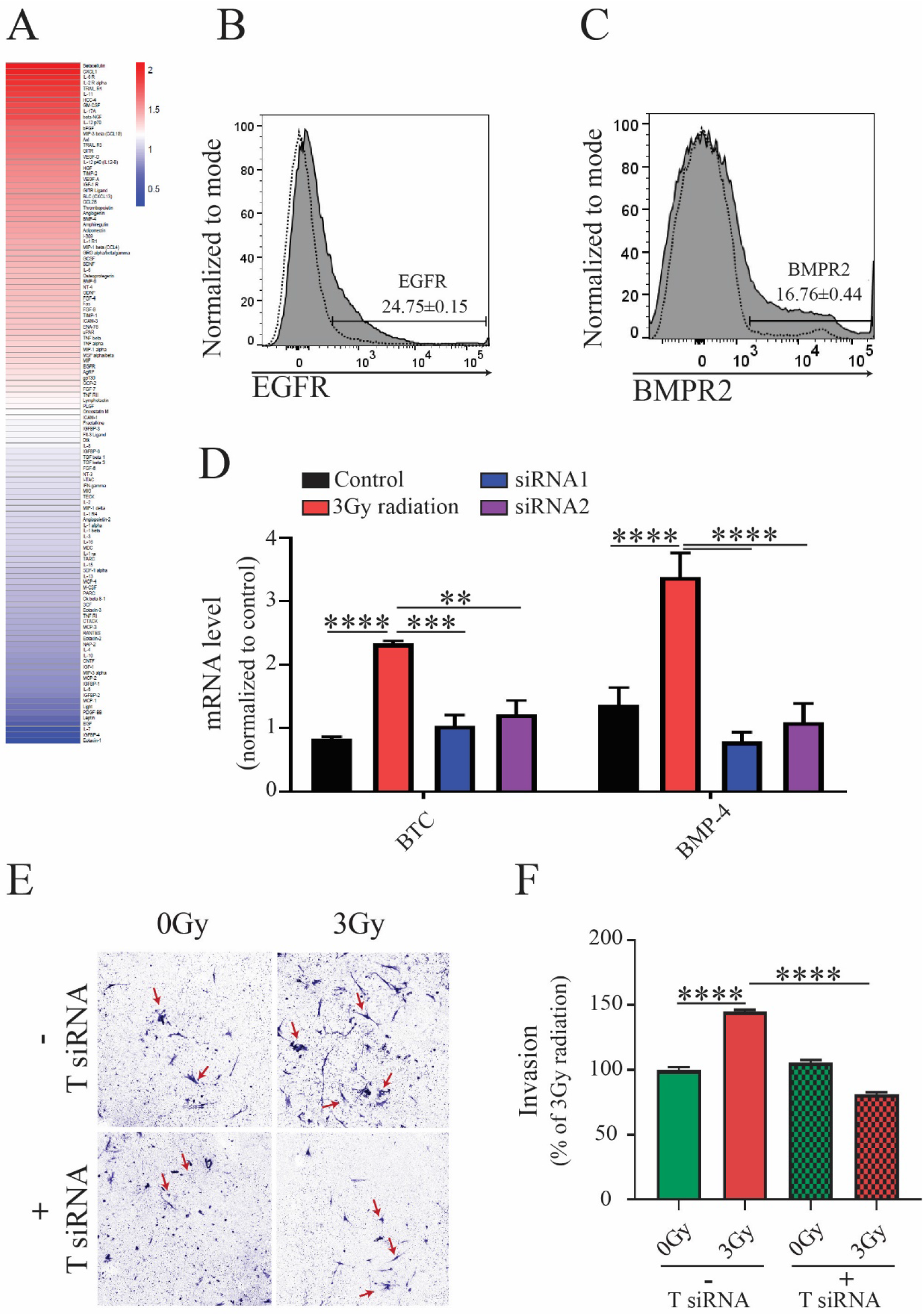
Radiation-induced upregulation of betacellulin (BTC) and bone morphogenic protein 4 (BMP-4) contributes to MSC migration to DIPG tumors. A) Protein array of SF8628 DIPG cells isolated from the brainstem of control and irradiated mice (n=3 per control and irradiated groups). Heat map values for each protein represent an average of two independent reads of the array. Representative flow cytometry histogram of MSCs for expression of B) EGFR-epidermal growth factor (n=3) and C) BMPR2-bone morphogenic protein receptor 2 (n=3), receptors for BTC and BMP-4 ligands, respectively. D) Quantitative PCR assessment of the mRNA levels of expression of BTC and BMP-4 after siRNA-induced gene silencing (n=3, One-way ANOVA with Tukey’s post-hoc test. Levels of significance are: ** p <0.01, *** p <0.001 and **** p<0.0001). E) Migration of MSCs toward irradiated SF8628 DIPG cells after silencing of both BTC and BMP-4 siRNA *in vitro*. E) Representative images of MSCs migrated through transwell membranes towards supernatants collected from control and 3Gy irradiated SF8628 DIPG cells. Red arrows indicate migrated cells. F) Quantification of migrated MSCs at the same conditions shown in E) (One-way ANOVA with Tukey’s post hoc test, n≤4. Level of significance: **** p<0.0001).

### OV infectivity, replication in, and cell killing of DIPG in vitro

We compared the efficacy of CRAd.S.pK7 and Ad5delta24RGD that have distinct modifications of their fibers that is necessary for cellular entry of OV. These viruses also differ with respect to the promoter that drives viral gene expression: survivin for CRAd.S.pK7, and CMV for Ad5delta24RGD. Both viruses successfully killed DIPG cells *in vitro* (Fig. 3 A-C), though CRAd.S.pK7 was more effective at a lower multiplicity of infection (MOI) in SF8628 and SF7761 cells (Fig. 3A, B). These findings are consistent with results showing superior binding of the CRAd.S.pK7 OV to the surface of the SF8628 over the SF7761 and DIPG007 cells (Fig. 3D).

**Figure 3.**
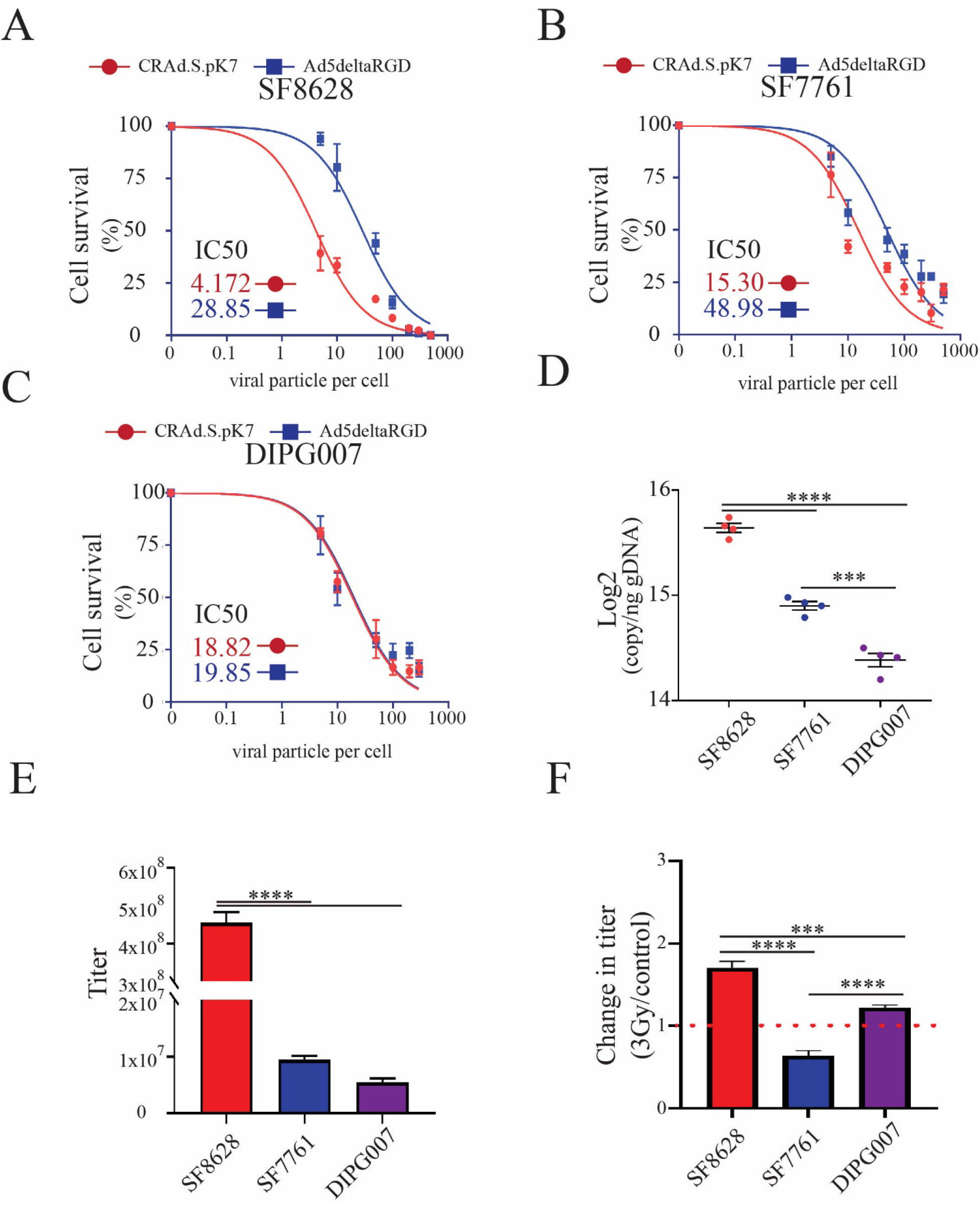
Oncolytic activity of CRAd.S.pK7 in patient-derived DIPG cells. A) Kinetics of killing by CRAd.S.pK7 and Ad5deltaRGD of SF8628 DIPG cells expressing firefly luciferase were measured by the loss of luciferase activity 3 days after treatment of cells with OV (n=4). B) Kinetics of killing by CRAd.S.pK7 and Ad5deltaRGD of SF7761 DIPG cells (n=4) and C) DIPG007 DIPG cells (n=4) were assessed as described in A. D) Binding assay of CRAd.S.pK7 with the surface of DIPG cell lines SF8628, SF7761, DIPG007 was performed to assess the difference in the the binding of viral partciles to the cell surface using PCR-based quantification of viral DNA (One-way ANOVA with Tukey’s post-hoc test, n=4. Levels of significance are: *** p <0.001 and **** p<0.0001). E) The release of CRAd.S.pK7 progeny from SF8628, SF7761, and DIPG007 DIPG cell lines was assessed by determining viral titer using staining for hexon viral protein in 293T cells (Negative binomial model was fitted and followed by Comparisons of Group Least Squares Means with Tukey-Kramer adjustment, n=4). F) Change in the progeny release of CRAd.S.pK7 from in DIPG cells after 3Gy irradiation treatment (One-way ANOVA with Tukey’s post-hoc test, n=4. Levels of significance are: * p <0.05, ** p <0.01, *** p <0.001 and **** p<0.0001).

Viral replication in SF8628 cells was 47.94±5.23 times higher than in SF7761 and 82.82±3.03 times higher than in DIPG007 cells (Fig. 3E); though all tested cells robustly generated and released viral progeny. In the context of irradiation, SF8628 and DIPG007 cells exposed to 3Gy irradiation produced 70.12±7.92% and 22.3±3.28%, respectively, more OV than non-irradiated control cells (p<0.0001), whereas a reduction by 44% in was observed in irradiated SF7761 (Fig. 3F).

To identify individual DIPG molecular characteristics that could help predict individual tumor responsiveness to CRAd.SpK7 therapy, both in the presence and absence of radiation, we used multiple approaches. The analysis of RNAseq data from DIPG tumor tissue^5^ revealed robust expression of mRNAs encoding integrin (SFig. 2A-C) and CAR (SFig. 2D), attachment proteins for virus. Analysis of patient DIPG tissue (Fig. 4A, B) and DIPG cell lines (Fig. 4C, D) showed robust expression of syndecan 1 and perlecan mRNAs, with syndecan 1 expression being highest in SF8628 cells, among all DIPG cell lines, as indicated by flow cytometry studies (Fig. 4E, F). Irradiation of SF8628 cells slightly reduced, by 6.67±0.02%, the presence of the syndecan 1 at the cell surface of treated cells (Fig. 4F).

**Figure 4.**
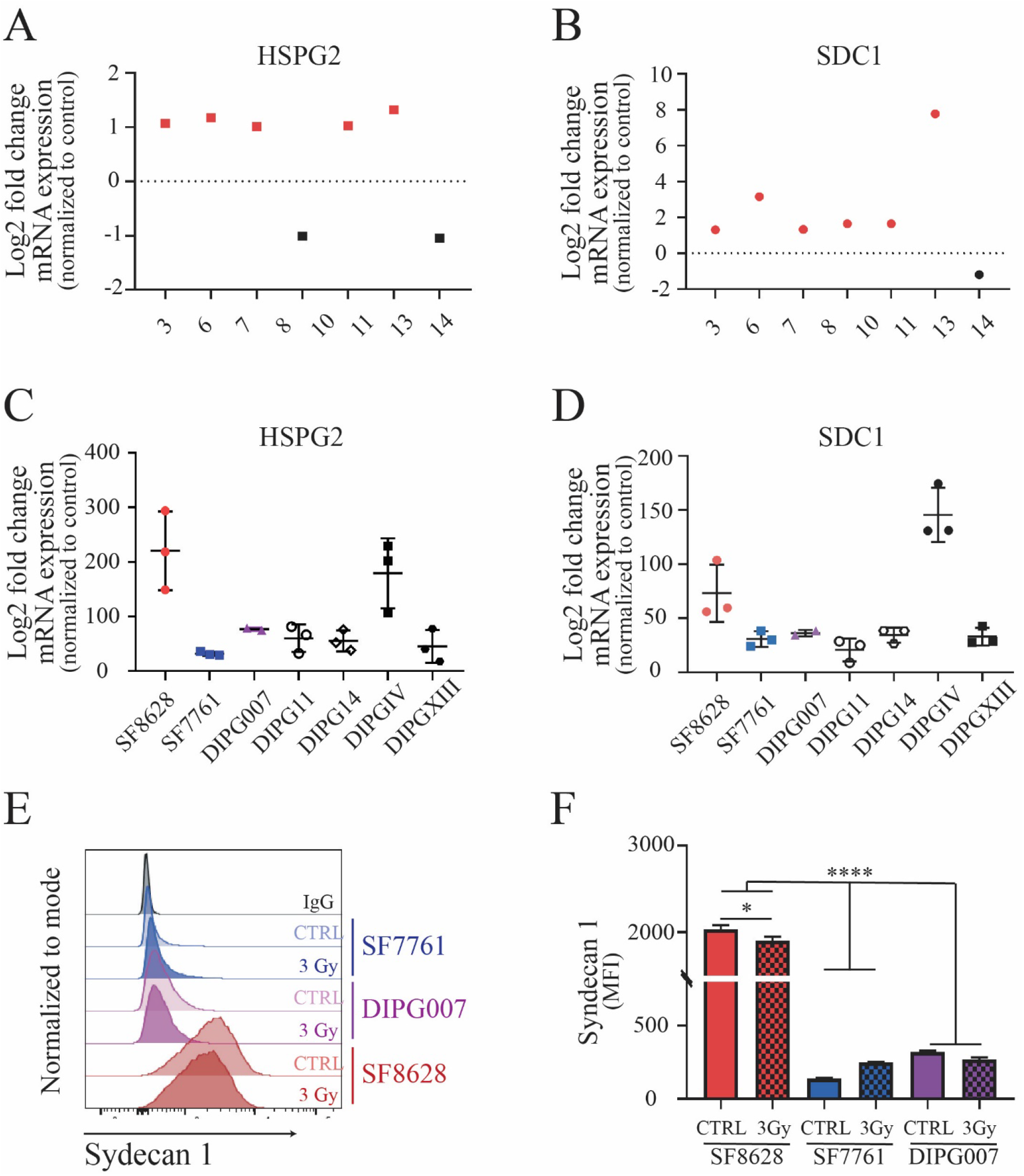
Expression of surface heparan sulfate proteoglycans in human DIPG tumor tissue, patient-derived cell lines, and mouse xenograft tumors. RNAseq analysis for the expression of A) heparan sulfate proteoglycan 2 (HSPG2) and B) syndecan 1(SDC1) in DIPG patient samples. RNAseq analysis of the patient-derived primary cell lines SF8628, SF7761, DIPG007, DIPG11, DIPG14, DIPGIV and DIPGXIII for expression of C) HSPG2 and D) SDC1. E) Representative flow cytometry histogram for the surface expression of syndecan 1 in DIPG cell lines SF8628, SF7761, and DIPG007 (n≥3) before and after fractionated irradiation treatment with 3Gy total dose. F) Analysis of syndecan 1 levels expressed on the cell surface in control and irradiation-treated DIPG cells detected by flow cytometry (Welch’s Anova test with Dunnett’s post hoc test, n=4-7. Levels of significance are: * p <0.05and **** p<0.0001.

As the survivin promoter drives the replication of CRAd.S.pK7 in cancer cells, we examined survivin expression *in silico* using RNA microarray data sets available at the GlioVis database^33^. This analysis showed that survivin, encoded by the *BIRC5* gene, is significantly overexpressed in a range of pediatric brain tumors as compared to non-tumor tissues (SFig. 3A, B). Analysis of RNAseq data set from patient tissues (Children’s Brain Tumor Tissue Consortium, CBTTC, https://cbttc.org/) revealed a significantly greater expression of survivin (<0.0001) in high-grade pediatric glioma compared to low-grade glioma (SFig. 3C)^34, 35^. Analysis of RNAseq data set^5^ showed a significant increase of survivin in the majority of human DIPG samples, when compared to normal brain tissue controls (Fig. 5A: p-value < 0.05). Analysis of RNAseq data obtained from the CBTTC (n=14) and the Pediatric Brain Tumor Atlas (PNOC003, n=31) also showed robust expression of *BIRC5* RNA in DIPG tissue (Fig. 5B, SFig. 4 A-B)^34, 35^. Immunostaining of DIPG tissues showed a varying degree of survivin expression among DIPG patients, in agreement with variability seen at the mRNA level (Fig. 5C). RNAseq and quantitative PCR analysis revealed robust *BIRC5* mRNA expression in all DIPG cell lines (Fig. 5D-E), and Western blot results showed high levels of survivin protein in SF8628 and DIPG007 cells (Fig. 5 F). We observed reduced survivin promoter activity, mRNA expression, and reduced protein levels in DIPG007 and SF7761 cells, but not in SF8628 irradiated cells (Fig. 6A-D). Consistent with the radiation-induced increase in survivin expression observed in SF8628 cells, combinatorial treatment with CRAd.S.pK7 and radiation increased the number of dying cells by an additional 24.78±2.65 % in comparison to control, after 5 days in culture (SFig. 5). These data indicate that CRAd.S.pK7 possesses potent lytic activity in DIPG cells with its replication being dependent on the expression of survivin and proteins mediating viral entry into DIPG cells.

**Figure 5.**
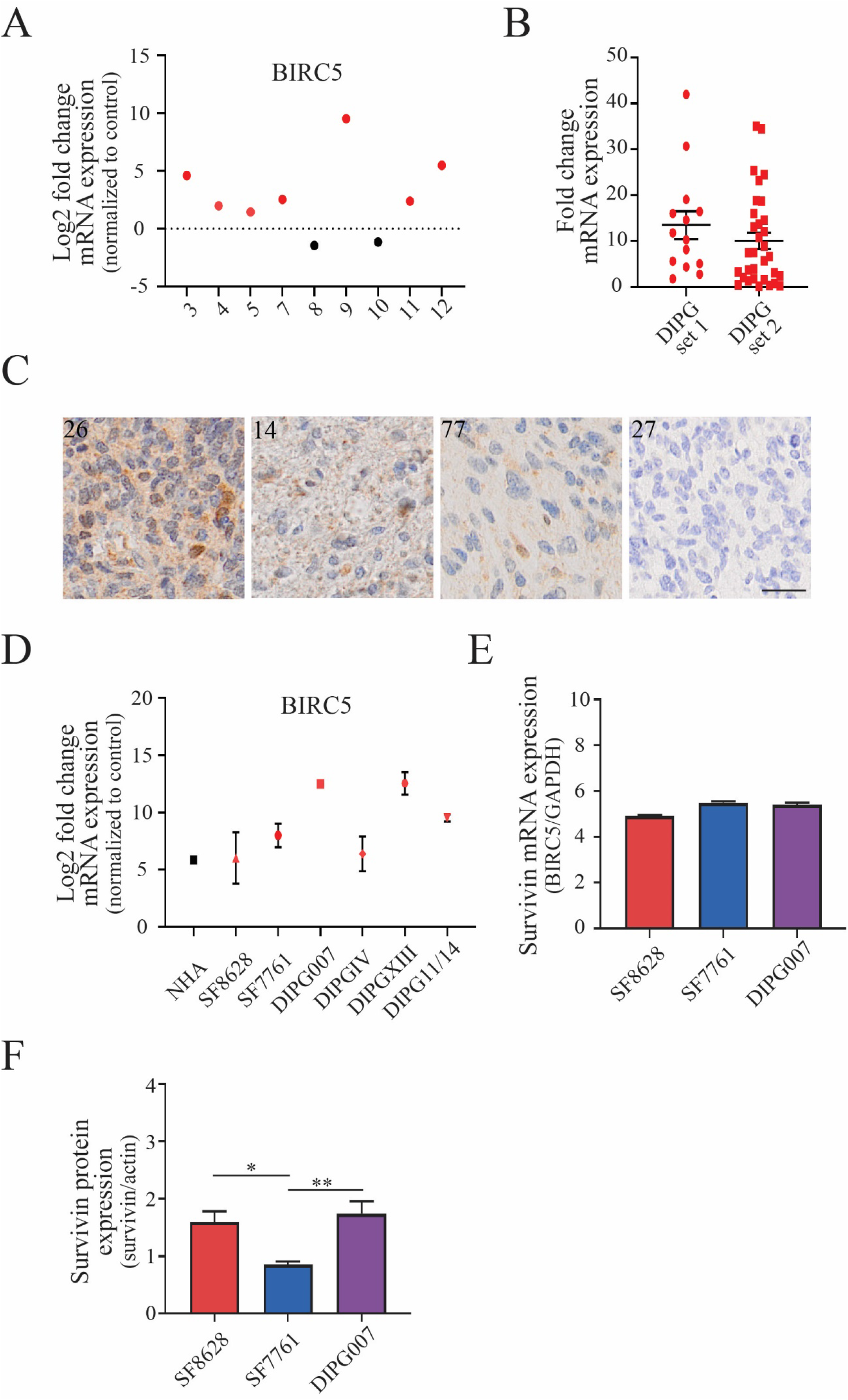
Survivin expression in human DIPG tumor tissue, patient-derived cell lines, and mouse xenograft tumors. A) Fold change in mRNA expression of the *BIRC5* gene encoding the survivin protein in human DIPG tumor tissue samples, normalized to average expression across normal brain tissue from the same patients. B) Fold change in *BIRC5* expression in human DIPG tumor tissue. ‘DIPG set 1’ data was acquired from the Children’s Brain Tumor Tissue Consortium (CBTTC) (n=14), and ‘DIPG set 2’ data was aquired from the Pediatric Brain Tumor Atlas (PNOC003) (n=31) dataset (PedcBioPortal). C) Immunohistochemitry of DIPG patient samples stained for survivin representing strong (score=III, patient 26), medium-strong (score=II, patient 14), weak (score=I, patient 77) and absent (score=0, patient number 27) expression (scale: 50μm). D) RNA expression of *BIRC5* gene in normal human astrocytes (NHA) and patient-derived DIPG cell lines. E) qPCR analysis of basal *BIRC5* mRNA expression in DIPG cell lines SF8628, SF7761, DIPG007. F) Western blot quantification of survivin protein expression normalized to actin expression in DIPG cell lines (One-way ANOVA with Tukey’s post-hoc test, n=3. Levels of significance are: * p <0.05 and ** p <0.01).

**Figure 6.**
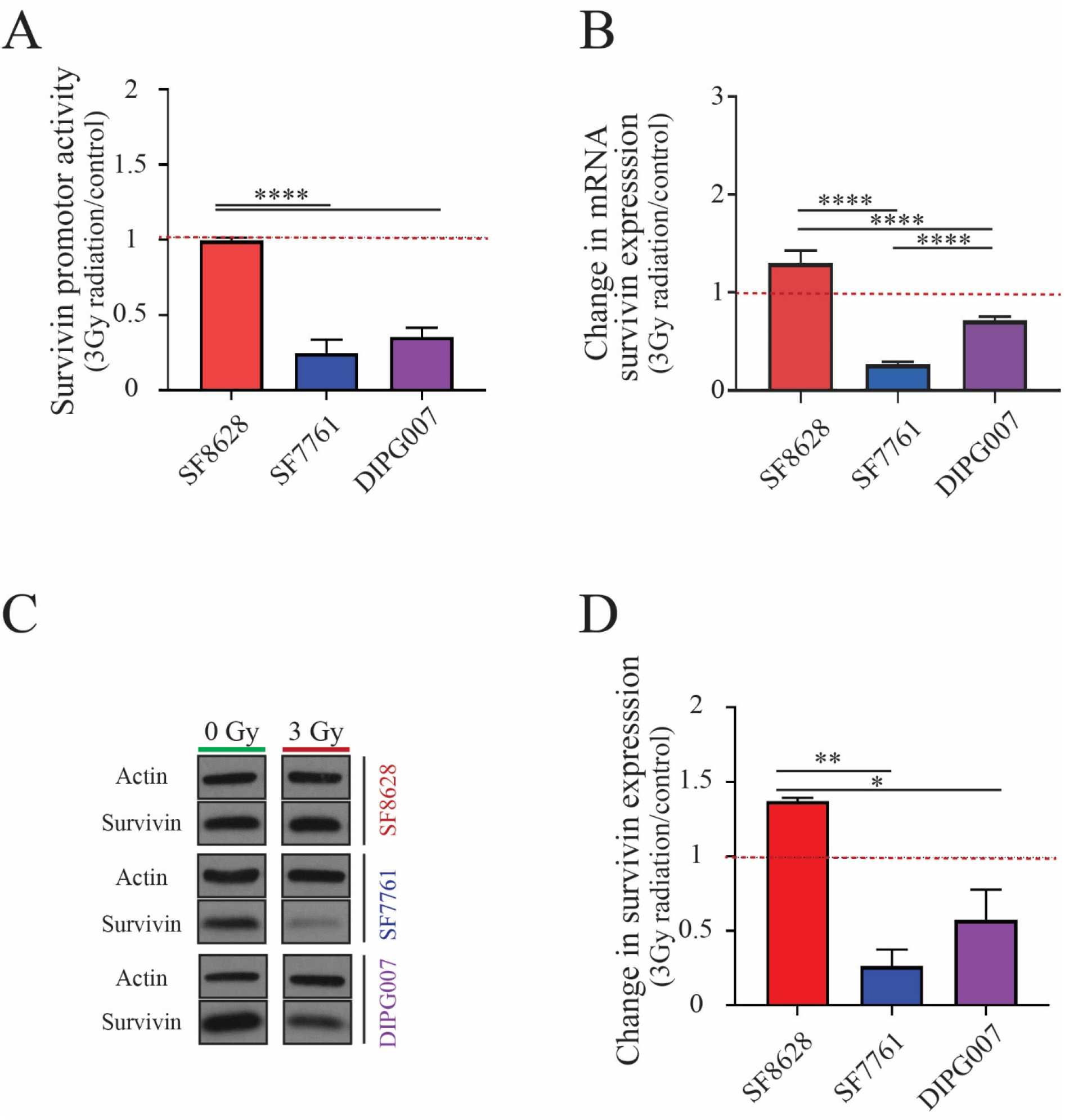
Expression of survivin is affected by irradiation in patient-derived cell lines. A) Survivin promoter activity in irradiated SF8628, SF7761, and DIPG007 cell lines normalized to untreated controls (One-way ANOVA with Tukey’s post-hoc test, n=6. Levels of significance: **** p<0.0001). B) qPCR analysis of surviving mRNA in response to irradiation (One-way ANOVA with Tukey’s post-hoc test, n=4. Levels of significance: **** p<0.0001). C) Example of survivin western blot of SF8628, SF7761, and DIPG007 DIPG cell lines before (0Gy) and after (3Gy) irradiation. D) Change in survivin protein expression in response to irradiation treatment (One-way ANOVA with Tukey’s post-hoc test, n=3. Levels of significance are: * p <0.05 and ** p <0.01).

### CRAd.S.pK7 treatment mediated by MSC delivery improves the survival of mice-bearing DIPG tumors, and survival benefit is enhanced when combined with fractionated irradiation

We performed a longitudinal analysis to establish the effect of irradiation on the tumor growth *in vivo*. Using BLI, we found that 3Gy radiation did not delay tumor growth of SF8628 DIPG PDX (Fig. 7A). Next, SF8628 mice bearing SF8628 brainstem xenografts were treated with CRAd.S.pK7-loaded MSCs following irradiation. There was no survival benefit observed in association with IND administration of cell-based OV (SFig. 6A), even though the expression of hexon, a viral capsid protein, was evident in the DIPG tissue (SFig. 6B). Local delivery of the MSCs-CRAd.S.pK7 to SF8628 DIPG xenograft tumors 2mm superficial to the tumor implantation site was not beneficial as a monotherapy (Fig.7B). However, the animal survival was significantly (p<0.0001) improved with MSCs-CRAd.S.pK7 treatment after tumor irradiation (Fig. 7C). Collectively, these data confirm the ability of MSCs to successfully deliver CRAd.S.pK7 to DIPG in the context of low dose fractionated irradiation, resulting in a robust therapeutic effect in mice bearing DIPG tumors.

**Figure 7.**
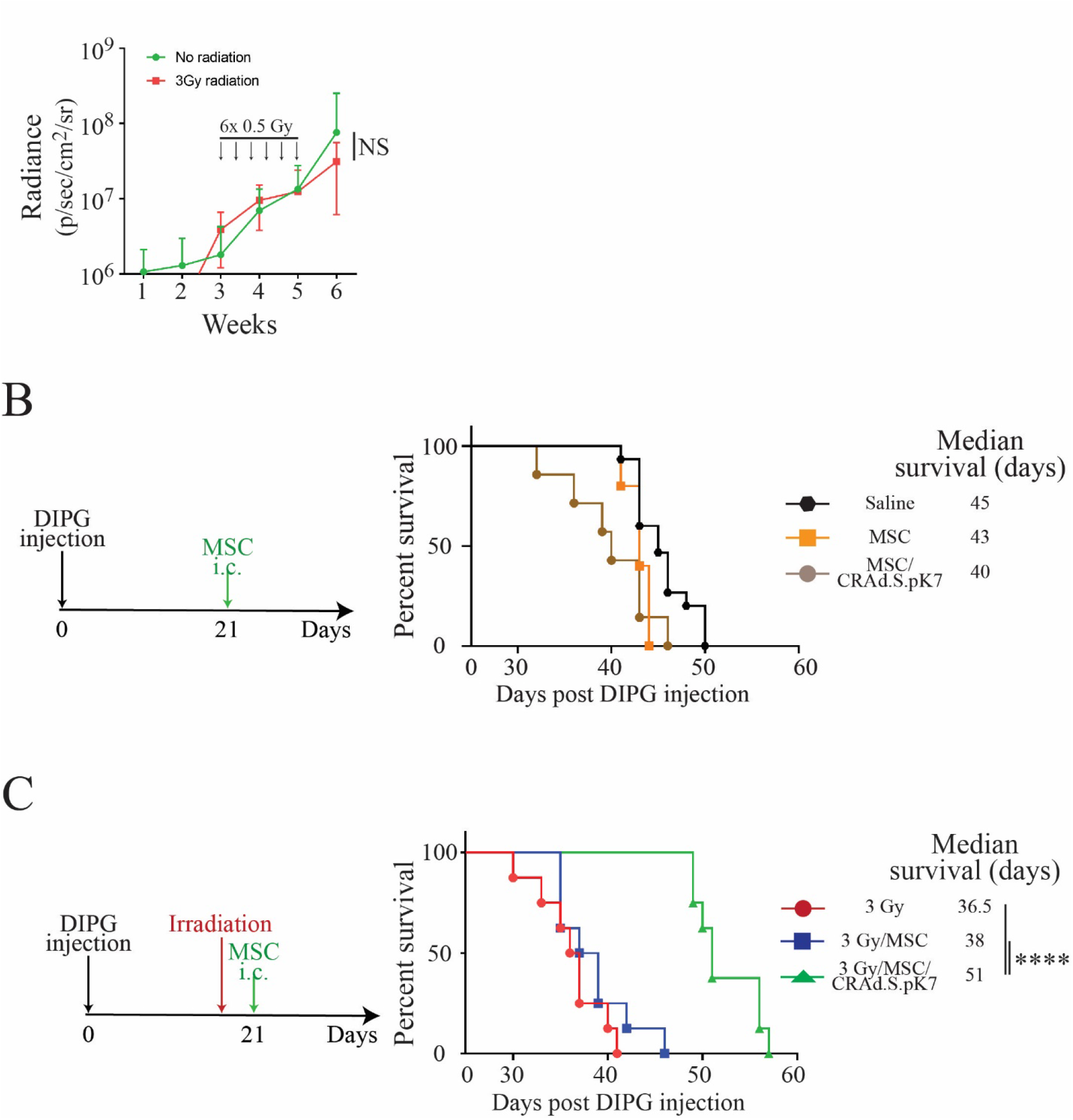
Survival analysis of mice treated with MSC-CRAd.S.pK7 via local versus intranasal delivery following radiation. A) Dynamics of growth of control and irradiated (total dose 3Gy) DIPG xenograft tumors were analyzed by serial BLI imaging. B) Animal treatment schema is shown on the left. Local delivery of 0.5×10^6^ /animal MSCs and MSC/CRAd.S.pK7 (Mentel-Cox Log Rank test, *p*=NS, N=15-saline, N=5-MSCs, N=7 MSC/CRAd.S.pK7). C) Animal treatment schema is shown on the left. Survival of DIPG bearing mice treated with locally delivered MSC/CRAd.S.pK7 after 3Gy fractionated irradiation (Kaplan-Meier survival curves were compared using log-rank tests with p-values adjusted with Bonferroni correction, n=8). Levels of significance **** p<0.0001).

## Discussion

Despite over 40 years of preclinical evaluation and clinical trials, DIPG persists as an incurable pediatric brain tumor and the number one cause of cancer-related death in children. CRAd.S.pK7 is a potent oncolytic agent presently being evaluated for safety in adult GBM patients (NCT03072134). In this study, we used patient-derived DIPG cell lines and mouse xenograft models^19, 36, 37^ to evaluate MSCs-loaded with the CRAd.S.pK7 for potential translation to the treatment of DIPG patients. Our analyses of patient tumor samples and patient-derived DIPG lines revealed the presence of cellular mediators necessary for the entry and replication of CRAd.S.pK7 in DIPG cells. Further, we have shown successful CRAd.S.pK7 replication in DIPG cells *in vitro*, and showed that low dose fractionated irradiation significantly improves animal survival when treated with MSCs carrying OV.

Delivery systems and platforms that overcome anatomical impediments posed by deep-seated brain tumors, such as DIPG, are in high demand. Stem cells, including MSCs, have low immunogenicity, can circumvent certain anatomic restrictions, such as the BBB, and navigate within the tight structure of the brainstem to home to the tumor site^38–41^. Neural and mesenchymal stem cells are well established for safe clinical use in the treatment of neurodegenerative and malignant diseases of the central nervous system, and can be used as carriers of therapeutic agents, including OVs^13, 14, 23, 30, 42, 43^. In the results presented here, we show that MSCs can deliver a therapeutic payload to brainstem xenografts in the context of fractionated irradiation^44, 45^. In fact, our results indicate that the irradiation of DIPG cells or DIPG-bearing mouse brains promotes MSC migration toward DIPG *in vitro* (SFig 1A) and *in vivo* (Fig. 1C, D). Analysis of cytokine and chemokine expression in untreated and irradiated DIPG tumors revealed a large number of upregulated factors capable of inducing cellular survival and stem cell homing to tumor (Fig. 2). Our results also show that the use of the survivin promoter to drive OV gene expression is appropriate in that the endogenous survivin gene, BIRC5, is highly expressed in most DIPG (Fig. 5).

OV-mediated tumor killing is also influenced by the ability of virus to infect tumor cells, and our analysis of patient-derived tumor tissue samples and cell lines showed that all DIPG express mRNA for viral attachment proteins syndecan 1 and perlecan, though at variable levels. High syndecan 1 and perlecan expression in the SF8628 DIPG cells was associated with higher viral progeny, relative to that of SF7761 and DIPG007 cells expressing lower levels of syndecan 1 and perlecan. Cumulatively, our results suggest that several tumor-specific molecular characteristics influence cell-based OV treatment efficacy.

Importantly, because radiation is the standard treatment for DIPG, we explored its effects on the context of IND and local delivery to the brain. The local, but not IND, delivery of MSC-OV resulted in significantly improved animal survival. This finding suggests sufficient quantity of oncolytic adenovirus must reach the tumor. Recent studies in preclinical models of DIPG and pediatric glioblastoma (GBM) demonstrated an encouraging efficacy of locally delivered OVs^46^. The upregulation of cytokine and chemokine expression in irradiated tumors suggests that modification of MSCs with corresponding receptors may also improve their migratory capacity and effective delivery of therapeutic load directly to the tumor site^29^.

CRAd.S.pK7 does not replicate in murine glioma cells, precluding its evaluation in an immunocompetent model. Martinez-Velez *et. al* recently showed that the efficacy of OV treatment in pre-clinical models of pediatric glioma depends on triggering the adaptive immune response^46^. These findings are in line with prior studies of multiple tumors types treated with OV that demonstrate a modulated immune response to OV administration that contributes to control of tumor growth^47–50^. However, because this immune response is also responsible for the rapid clearance of viral particles, a delay in OV elimination could improve the therapeutic efficacy of this anti-cancer treatment strategy^51–55^. In that respect, OV delivery using MSCs has been shown to protect OV from rapid viral clearance by the immune system, and consequently to extend the window of oncolytic activity within the tumor, increasing therapeutic efficacy^14^

In conclusion, our study supports oncolytic virus CRAd.S.pK7 encapsulated within MSCs, when used in combination with low dose radiation, as a therapeutic strategy that merits further investigation and potential translation for DIPG treatment.

## Materials and Methods

### Patients specimens

Patients specimens were collected at postmortem in accordance with Children’s National Medical Center with IRB approval (IRB #Pro00001339) as described previously^5^.

### Cell lines

Primary SF8628, SF7761, and DIPG007 were used. Detailed information can be found in supplemental materials.

### Bioluminescent imaging and MRI

Tumor visualization using MRI was done according to a previously described protocol^22^ Detailed information on BLI can be found in supplemental materials.

### Radiation Treatment

Experimental mice were exposed to 0.5 Gy radiation 3 times a week for 2 weeks to reach a total dose of 3 Gy. Detailed procedures are described in materials and methods.

### MSC labeling

Detailed information on procedure can be found in supplemental materials.

### Histology

Detailed procedures can be found in supplemental materials.

### Quantitative analysis of MSCs migration in vivo

Details are described in supplemental materials.

### Immunocytochemistry

Details of procedure are described in supplemental materials.

### Western Blot

Details of procedure are described in supplemental materials.

### Quantitative PCR

Procedure described in supplemental material

### Invasion Assay

Procedure described in supplemental material

### Cytokine Array

Procedure described in supplemental material

### siRNA silencing

Details are described in supplemental materials.

### Flow cytometry

Flow cytometry was performed on the BD Fortessa flow cytometer (Becton Dickinson, Franklin Lakes, NJ) at the Robert H. Lurie Comprehensive Cancer Center Flow Cytometry Core Facility. Detailed procedures can be found in supplemental materials.

### Quantitative analysis of the binding CRAd.S.pK7 to the cell surface

Details are described in supplemental materials.

### Evaluation of CRAd.S.pK7 replication and toxicity

To assess the release of viral progeny, DIPG cells expressing firefly luciferase were infected with CRAd.S.pK7 at MOI 10 v.p./cell in DMEM media supplemented with 1%FBS. Detailed procedures for assessment can be found in supplemental materials.

### Analysis of Survivin Promoter Activity in DIPG Cells

To assess the survivin promoter activity, we employed a replication-deficient adenoviral vector expressing firefly luciferase under the control of the survivin promoter as previously described^56^. Detailed procedures for assessment can be found in supplemental materials.

### Animals and Surgical Procedures

Six-week-old female athymic mice (BALB/c background) were purchased from Charles River and housed under aseptic conditions. Pontine injection of tumor cells was performed as described previously^37^. Detailed surgical procedures are described in supplemental materials.

### Intranasal Delivery

Mice were treated with methimazole (50 mg/kg body weight) 2 days before IN-MSC delivery as previously described^22^. Detailed infromations can be found in supplemental materials

### Survival Analysis

Mice with similar tumor size (as quantified using BLI) were treated with therapeutic MSCs loaded with oncolytic adenovirus. MSCs bearing oncolytic adenovirus were delivered using either through IN route or intracranial injection into the brain at a location proximal to the DIPG tumor at a depth of 3 mm.

### Statistics and data analysis

The analyses of collected data were performed using GraphPad 8 Software (Prism, La Jolla, CA) and SAS 9.4 (SAS Institute Inc., Cary, NC). RNAseq was analyzed as previously published^57, 58^. Analysis of RNAseq from The Children’s Brain Tumor Tissue Consortium (CBTTC) and Pediatric Brain Tumor Atlas (PNOC003) obtained from PedcBioPortal (https://pedcbioportal.kidsfirstdrc.org/) was previously described^34, 35^. Details on statistics and data analysis can be found in supplemental materials.

## Acknowledgments

This work was supported by NIH R01NS087990 and P50CA221747 SPORE for Translational Approaches to Brain Cancer. We are also grateful to the Nora Redman Endowment Fund of the Community Foundation of Louisville, and to the Isabella Kerr Molina Foundation for generous gifts to support the study. Some of the materials employed in this work were provided by the Texas A&M Health Science Center College of Medicine Institute for Regenerative Medicine at Scott & White through a grant from ORIP of the NIH, Grant # P40OD011050. This work was also supported by the Northwestern University – Flow Cytometry Core Facility supported by Cancer Center Support Grant (NCI CA060553) and the Center for Advanced Microscopy generously supported by NCI CCSG P30 CA060553 awarded to the Robert H Lurie Comprehensive Cancer Center.

## Author contributions

MC, KP, RH, HK, LI, YH, EB, CH, AS, and IB designed and performed experiments. KP, TX performed statistical analysis. MC, KP, ML, RH, AS, CDJ, DC, JN and IB contributed to the manuscript writing.

