## Supplemental Materials for "MSCs Successfully Deliver Oncolytic Virotherapy to Diffuse Intrinsic Pontine Glioma"

**Supplemental Methods**

*Cell lines*

Primary SF8628, SF7761, and DIPG007 were used. (H3.3K27M DIPG) cell lines were generously provided by Dr. Rintaro Hashizume (Northwestern University). DNA fingerprints, using the Powerplex16HS System (Promega DC2101), were obtained to confirm the identity of the cell lines. The establishment of SF8628 from a surgical specimen, and tumor cell modification for expression of firefly luciferase, for *in vivo* bioluminescence imaging, has been described by Hashizume et al.[37]. DIPG007 (H3.3K27M DIPG) cell line was kindly provided by Dr. Angel Montero Carcaboso (Hospital Sant Joan de Déu, Barcelona, Spain). Human bone marrow-derived MSCs were purchased from the Texas A&M Health Science Center College of Medicine Institute for Regenerative Medicine and were grown in MEMalpha supplemented with 10% FBS.

*Bioluminescent imaging and MRI*

To visualize DIPG tumors in animals, mice were injected intraperitoneally (i.p.) with 200 µl (4.5 mg per animal) D-luciferin sodium salt (Gold Biotechnology, St Louis, MO). Signal was recorded as photon counts using Caliper Life Sciences IVIS Spectrum and analyzed with Living Image 3.1 software.

*Radiation Treatment*

Experimental mice were exposed to 0.5 Gy radiation 3 times a week for 2 weeks to reach a total dose of 3 Gy. The entire body of the animals, excluding the head, was covered with lead shields. Mice were anesthetized with ketamine/xylazine mixture for the duration of radiation exposure. DIPG cells *in vitro* were exposed to 1, 2, or 4 Gy radiation each day for 3 days, 24 h after initial plating to reach a total dose of 3, 6, and 12 Gy, respectively. Cells were analyzed after recovering from treatment for 1, 2, or 3 days.

*MSC labeling*

MSCs were labeled with superparamagnetic iron oxide particles (SPIOs) (Bangs Laboratories, Fishers, IN) at 20 particles per cell in the OptiMEM culture medium for 6 hours. MEMalpha media with 10% FBS was then added to the plate for overnight incubation. The efficiency of MSC labeling with SPIOs was determined by Prussian Blue staining. Labeled cells were suspended in PBS in a volume of 12 μl/animal for IND.

*Histology*

Tumor morphology was assessed using H&E solutions from Sigma Aldrich (Hematoxylin MHS32, Eosin HTT10232). For Prussian blue staining, frozen 10 μm brain sections were fixed in 4% paraformaldehyde and, after three washes with PBS, stained with a 1:1 solution of 10% potassium hexacyanoferrate trihydrate with 20% HCl (Sigma-Aldrich, St Louis, MO) and counterstained with nuclear Fast Red stain (Vector Laboratories, Burlingame, CA). Tumor morphology was assessed using Hematoxylin and Eosin staining. DIPG and MSCs were detected by immunofluorescence and analyzed using a Leica Microsystems microscope.

Paraffin-embedded sections (5μm) from patients' DIPG tissues were de-paraffinized, and antigen-retrieval was performed using 10 mM sodium citrate (pH 6). Primary anti-survivin antibody (Abcam 60.11) was used for staining at dilution of 1:500. After overnight incubation at $4^{0}$C, slides were probed with anti-mouse secondary antibodies at 1:200 dilution. Antibody binding was visualized using 3,3'-diaminobenzidine (DAB) detection with Leica BOND-MAX automated strainer (Leica Biosystems). Slides were imaged at 20X magnification. Staining was scored by an experienced neuropathologist.

*Quantitative analysis of MSCs migration in vivo*

Tissue sections were collected from the brain of mice treated with SPIO-labeled MSCs by intranasal delivery and stained for Prussian blue. Each experimental group consisted of 3 animals. Ten slides (with 2-3 tissue sections on each slide collected from the ventral side of the brainstem, 150µm apart each other) were imaged in the Cytation 5 instrument (Biotek, Winooski, VT) at a magnification of 20X.

*Quantitative analysis of MSCs migration in vivo*

Images were stitched together using the Gen5 software to generate a composite image of the brainstem. Images were processed using ImageJ through color separation (RGB) and isolation for blue clusters. Clusters were differentiated from the background signals through the deconvolution function of the Image J software. Prussian blue quantification was calculated in Image J software by counting the intensity of the blue signal.

*Immunocytochemistry*

MSCs were detected using AlexaFluor 488-conjugated primary antibody specific for CD105 (Life Technologies). DIPG cancer cells were detected using a rabbit primary antibody for H3 K27M (Millipore) and a Cy3-conjugated anti-rabbit secondary antibody (Invitrogen). To evaluate the intratumoral delivery of viruses by MSC, tumor-bearing animals were treated via IND or local delivery with MSCs-CRAd.S.pK7. Five days later, animals were euthanized, brain tissue collected, snap-frozen, and sectioned. Tissue sections were stained for the expression of hexon viral protein as it has been previously described by Spencer et al.

*Western Blot*

Protein lysates were run on a 4-20% PAGE and transferred to the PVDF membrane using wet or semi-dry transfer. The Tris-HCl (4-20%) gel and PVDF membrane, ImmunStar WesternC, were purchased from Bio-Rad, (Hercules, CA). Antibodies for GAPDH (Cell Signaling 14C10) and

Survivin (Novus NB500-201) were used. Secondary antibodies were purchased from Santa Cruz Biotechnology (sc-2357) and Jackson labs, Biotin SP-conjugated donkey anti-rabbit IgG (711-065-152), and Peroxidase-conjugated Streptavidin (016-030-084).

*Quantitative PCR*

DIPG cells were harvested for RNA using the Qiagen RNeasy kit, and cDNA was generated using the BioRad iScript kit. Primers were designed to detect survivin and the top-expressing cytokines for qPCR analysis. GAPDH-F: 5'-GGTCGGAGTCAACGGATTTGG-3', GAPDH-R: 5'-CATGGGTGGAATCATATTGGAAC-3', Survivin-F: 5'-ACCACCGCATCTCTAC-3', Survivin-R: 5'-TCCTCTATGGGGTCGT-3', BTC-F 5'-AATTCTCCACTGTGTGGTAGCA-3', BTC-R: 5'-GGTTTTCACTTTCTGTCTAGGGG-3', BMP4-F: 5'-TTCCTGGTAACCGAATGCTGA-3', BMP4-R: 5'-CCCTGAATCTCGGCGACTTTT-3'

*Invasion Assay*

The invasion was measured using Corning BioCoat Matrigel Invasion Chambers. DIPG cells were plated on the bottom wells in DMEM with 1% FBS and treated with irradiation or siRNAs. MSCs were plated at a density of $10^4$ cells in the top chamber in 1% FBS. Cells were fixed and stained 24 hours later for counting.

*Cytokine Array*

DIPG cytokine expression was performed using a cytokine antibody array purchased from Ray Biotech (AAH-CYT-1000).

*siRNA silencing*

DIPG cells were transfected with siRNA using MISSION siRNA Transfection Reagent (Sigma S1452). 6 µl of Transfection Reagent was combined with 12 pmol of siRNA in 200 µl of serum-free media for 10 minutes. The transfection mix was added to 200,000 DIPG cells in a 6-well plate and incubated overnight. siRNA constructs: BTC (NM_001729: SASI_Hs01_00219907, SASI_Hs01_00219908), BMP4 (NM_001202: SASI_Hs01_00122023, SASI_Hs01_00122020).

*Flow cytometry*

MSCs dissociated to the single-cell suspension were stained with fixable viability dye (APC/Cy7, Thermo Fisher), blocked with PBS/2% fetal bovine serum (FBS) and Human Fc blocking reagent (BioLegend, San Diego, CA). Following the Fc blockade, anti-EGFR PE-conjugated antibodies (BioLegend, San Diego, CA) or anti-BMPR2 monoclonal unconjugated antibodies (Thermo-Fisher) were added and incubated on ice for 30 min. Detection of anti-BMPR2 antibodies was done using FITC-conjugated donkey anti-mouse IgG after 30 min incubation on ice. Anti-syndecan 1 PE-conjugated antibodies were purchased from BioLegend. The analysis was performed using FlowJo 8 (Becton Dickinson, Franklin Lakes, NJ) analysis software.

*Quantitative analysis of the binding CRAd.S.pK7 to the cell surface.*

To assess the binding of OV with different human DIPG lines, cells were seeded at $4x10^5$ onto 6-well plates in DMEM media supplemented with 10%FBS. The next day, media was replaced with DMEM media supplemented with 1%FBS, the cells were pre-chilled on ice and then incubated with CRAd.S.pK7 at MOI 1 v.p/cell on ice for 60 min. The cells were rinsed with ice-cold PBS and collected for isolation of genomic DNA using the DNeasy Blood and Tissue kit (Qiagen, Valencia, CA). The polymerase chain reaction (PCR) amplification of adenoviral genomic DNA was performed using the Adeno-X™ qPCR Titration Kit (Takara). All data were normalized per ng of genomic DNA.

*Evaluation of CRAd.S.pK7 replication and toxicity*

To assess the release of viral progeny, DIPG cells expressing firefly luciferase were infected with CRAd.S.pK7 at MOI 10 v.p./cell in DMEM media supplemented with 1%FBS. Three days later, the supernatant of cells was collected and titrated over HEK 293 cells. In 48 hours, cells were washed, fixed and stained for the expression of viral protein, hexon, using Adeno-X™ Rapid Titer Kit (Takara). The number of infectious units in each well was calculated according to the manufacturer's protocol. To assess the lytic activity of CRAd.S.pK7 and CRAd.delta24.RGD, DIPG cells plated at $5x10^3$ cells/well in 96 well plates were infected at various MOI. Seventy-two hours later, the luciferase activity in surviving cells was determined using the ONE-Glo™ Luciferase Assay System (Promega, Madison, WI) according to the manufacturer's protocol.

*Analysis of Survivin Promoter Activity in DIPG Cells*

DIPG cells were infected at MOI 10 in DMEM media supplemented with 1% FBS. Twenty-four hours later, cells were lysed in passive lysis buffer, and luciferase activity was determined using the ONE-Glo™ Luciferase Assay System (Promega, Madison, WI) according to the manufacturer's protocol. Experiments were performed in triplicate, and luciferase activities are presented as relative light units (RLU) normalized to the protein.

*Animals and Surgical Procedures*

Six-week-old female athymic mice (BALB/c background) were purchased from Charles River and housed under aseptic conditions. Pontine injection of tumor cells was performed as described previously Hashizume et al. Briefly, each mouse was injected with 1 μL of a cell suspension (10,000 cells/μL) into the pontine tegmentum at 1.5 mm to the right of the midline, posterior to the lambdoid suture, and at a depth of 5 mm from the inner base of the skull. A freehand technique was applied using a 26-gauge Hamilton syringe (Hamilton Company). All procedures were carried out under sterile conditions. All animal protocols were approved by the Northwestern University

Institutional Animal Care and Use Committee. Local delivery of MSCs was performed in the same injection site at a depth of 3 mm.

*Intranasal Delivery*

For IN delivery, mice anesthetized with isoflurane and given three treatments of 4 µl of $5x10^5$ MSCs in the nasal bed at 5-minute intervals. The procedure was carried out four times with 1-week interval.

*Statistics and data analysis*

In all experiments, significance was defined as the *p*-value of less than 0.05. Count data were compared by fitting Poisson regression model or negative binomial regression model as appropriate with Tukey-Kramer's adjustment for multiple comparisons. Continuous variables were analyzed using student's T-test or Welch's T-test for two groups, one-way ANOVA with Tukey's post-hoc test or Welch's test with Dunnett's post-hoc test for multiple groups as appropriate, and two-way ANOVA with Tukey's post-hoc test for groups with two independent variables. Survival curves were graphed via Kaplan-Meier method, and compared by Log-rank test or Renyi test, as appropriate, and p-values were adjusted using Bonferroni correction. Data are presented as mean ±SEM. Human tissue mRNA sequencing and proteomic analysis data were generated and analyzed as previously published, using Partek Genomics Suite v6.6 (Partek Incorporated, St. Louis, MO) [5]. Protein and gene expression values in tumor tissue were normalized and compared to control specimens from the same patient: *p-value* <0.05 and $\log_2$ fold change (FC) of expression ≤-2 or ≥2 compared to normal tissue were considered significant (1-way ANOVA). For patients from whom normal tissue was not available, comparisons were conducted using normalized values for all tumor and normal specimens and filtered for significance.

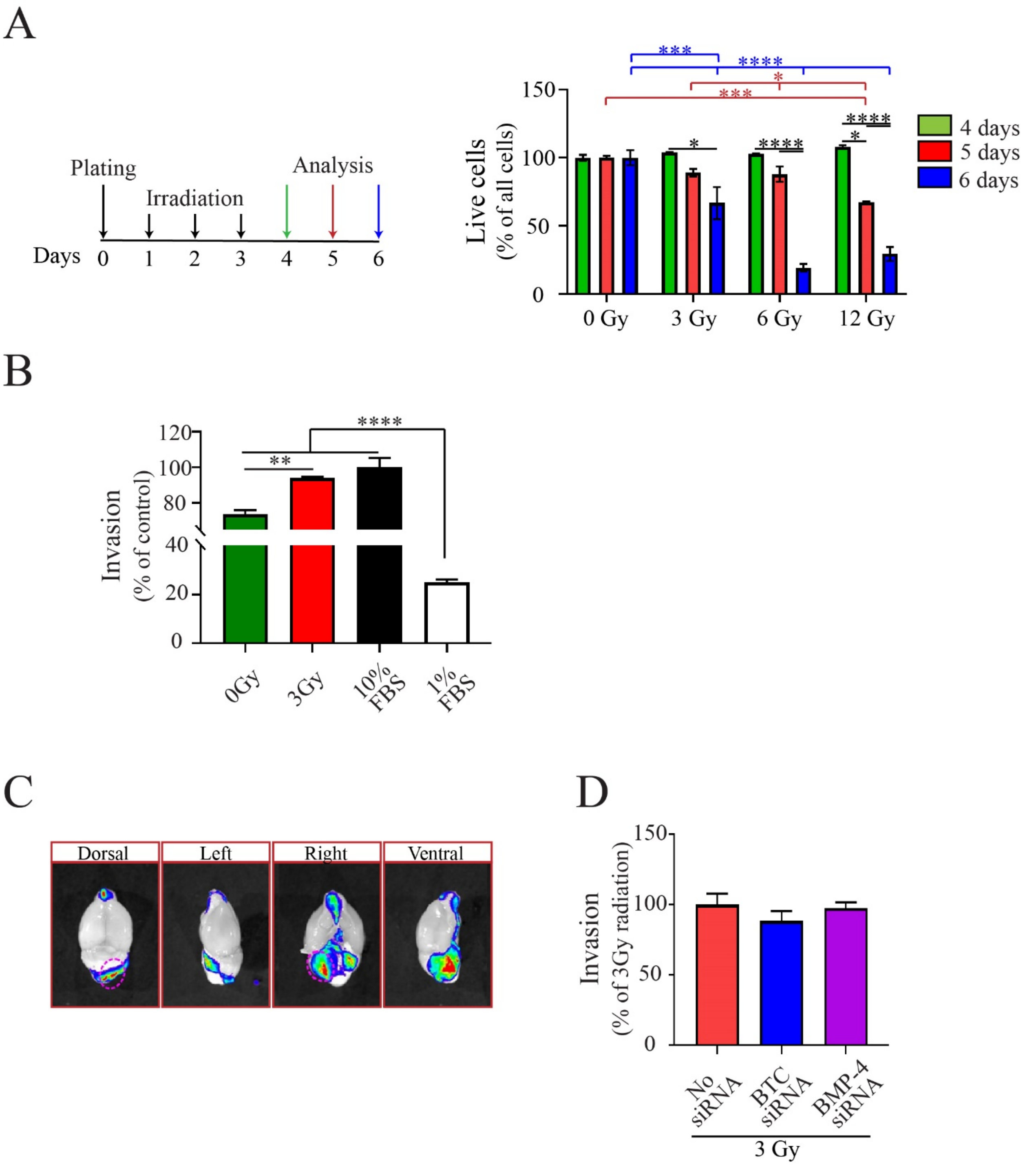


**Supplemental Figure 1. Effects of radiation on the viability of DIPG cells.** A) SF8628 human DIPG PDX cells were irradiated daily in 3 fractions for a total dose of 0, 3, 6, or 12 Gy, and analyzed on day 4, 5, and 6 in culture for their viability (Two-way ANOVA with Tukey's post-hoc test. Levels of significance: *p≤0.01, ***p≤0.0001, **** p<0.0001). B) MSCs demonstrated enhanced migration toward condition media of SF8628 DIPG cells treated with 3Gy irradiation

(One-way ANOVA with Tukey's post-hoc test, n=3. Levels of significance: **$p \leq 0.01$, ***$p \leq 0.0001$, **** $p<0.0001$). C). Representative image of firefly luciferase-expressing SF8628 DIPG tumors in the mouse brain. DIPG tissue was excised from the mouse brain using the imaging guidance and processed for analysis of cytokines/chemokines expression in the protein array. D) Migration of MSCs toward irradiated SF8628 DIPG cells after silencing of BTC or BMP-4 *in vitro*.

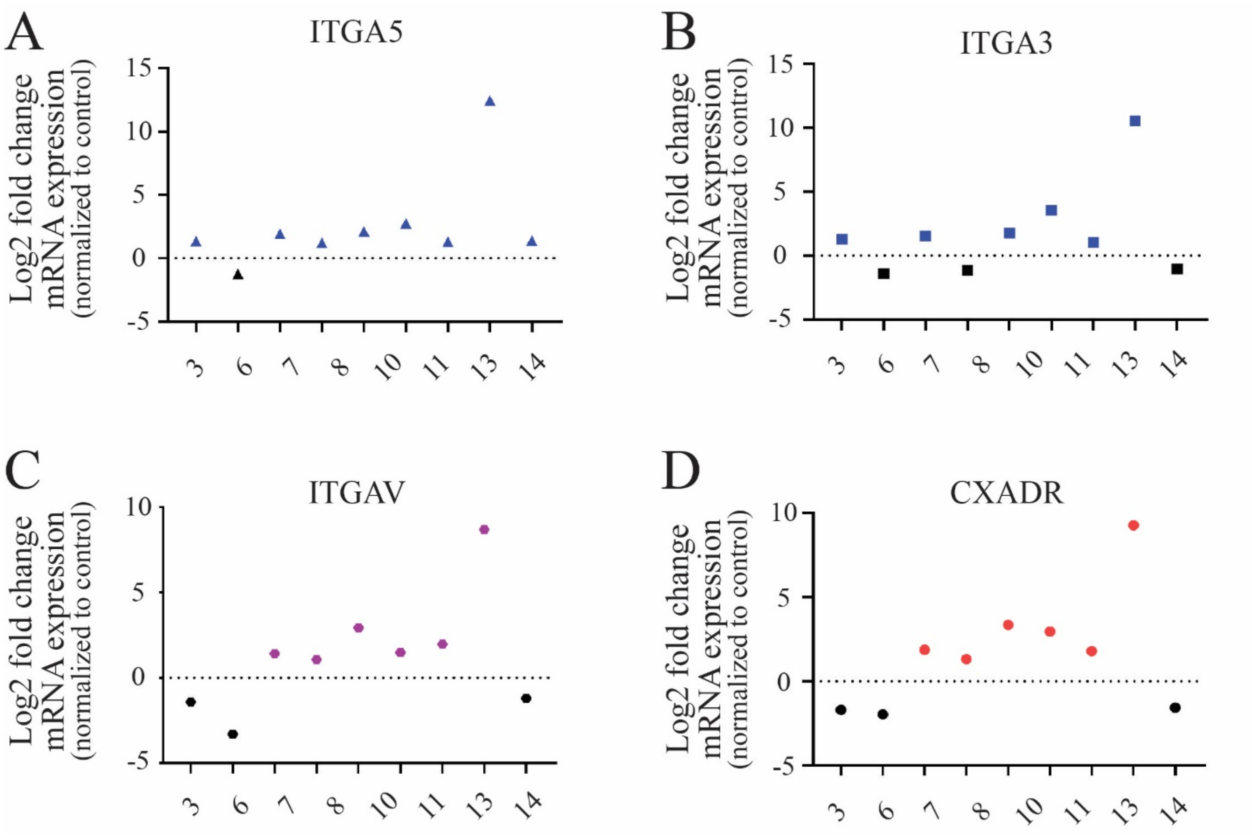


**Supplemental Figure 2. Expression of adenoviral entry receptors in patient DIPG tissue, patient-derived cell lines, and mouse xenograft tumors.** RNAseq analysis for the expression of A) integrin subunit alpha 5 (ITGA5); B) integrin subunit alpha 3 (ITGA3); C) subunit alpha V (ITGAV) and D) coxsackie virus and adenovirus receptor-CAR (CXADR) in human DIPG tumor tissue.

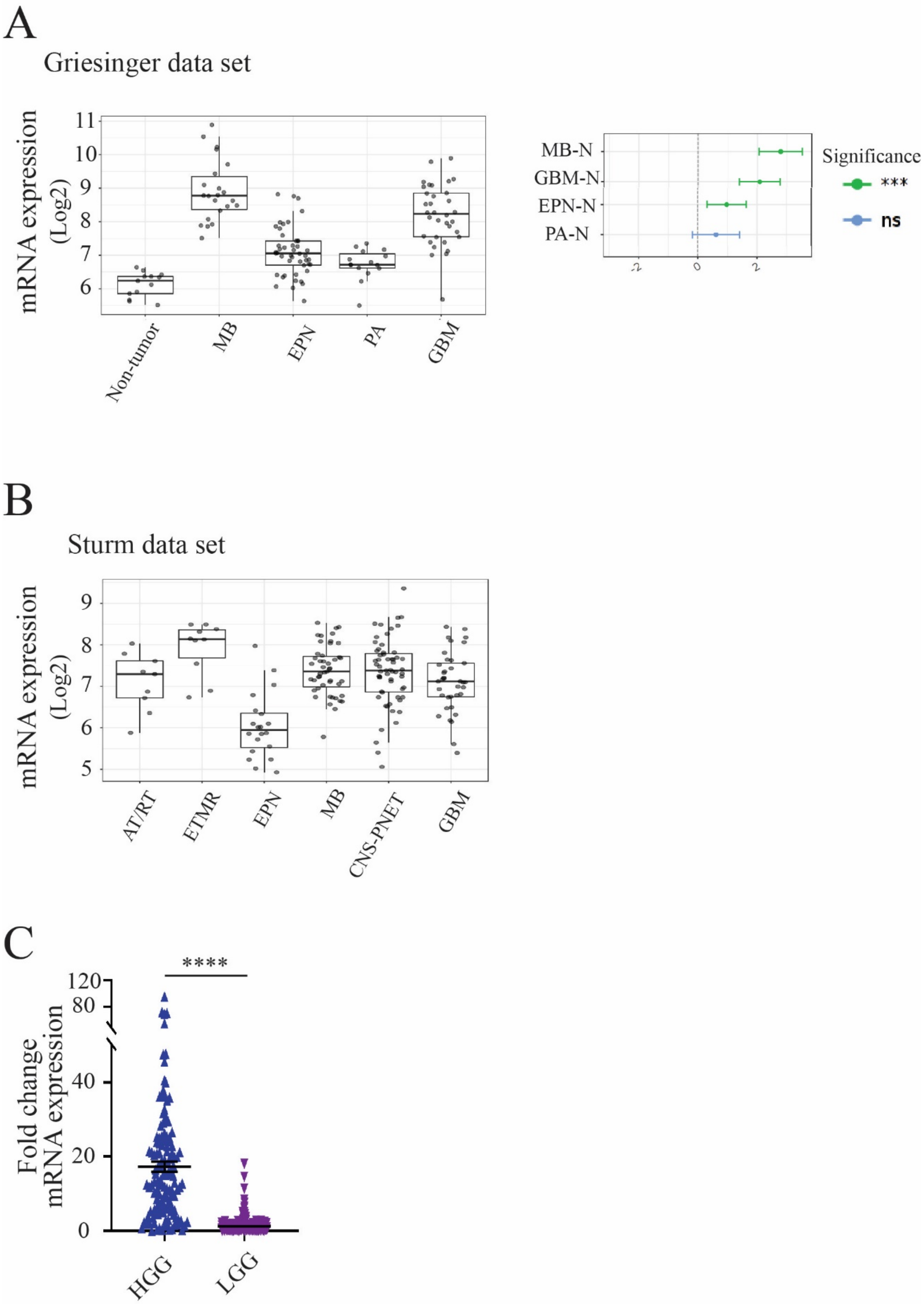
A
Griesinger data set
mRNA expression (Log2)
Non-tumor
MB
EPN
PA
GBM
MB-N
GBM-N
EPN-N
PA-N
Significance
***
ns
B
Sturm data set
mRNA expression (Log2)
AT/RT
ETMR
EPN
MB
CNS-PNET
GBM
C
****
Fold change mRNA expression
HGG
LGG

**Supplemental Figure 3. Analysis of survivin expression in pediatric brain cancers.** *In silico* analysis of A) the Grisinger data set was performed using the GlioVis portal. MB-medulloblastoma, GBM-pediatric glioblastoma, EPN- ependymoma, PA- pilocytic astrocytoma. Statistical significance was determined via the pairwise T-test using the GlioVis portal. B) The study of the Sturm data set was also performed using the GlioVis portal. N-normal tissue, AT/RT-atypically teratoid rhabdoid tumor, ETMR- embryonal tumors with multilayered rosettes, EPN-ependymoma, MB-medulloblastoma, GBM-pediatric glioblastoma, CNS-PNET- Central nervous system primitive neuroectodermal tumors. C) Fold change in *BIRC5* mRNA expression in HGG-high grade glioma (n=132) and LGG-low grade glioma (n=254) tumor tissue acquired from the Children's Brain Tumor Tissue Consortium (CBTTC) (Welch's T test. Level of significance: ****$p<0.0001$).

A

| Numerical order | SAMPLE_ID | Z score |
|---|---|---|
| 1 | 7316-549-T-332057 | 0.307202211 |
| 2 | 7316-1708-T-362697 | -0.416124525 |
| 3 | 7316-715-T-339877 | 1.859196219 |
| 4 | 7316-719-T-339937 | -0.150643284 |
| 5 | 7316-1695-T-362502 | 1.606983357 |
| 6 | 7316-639-T-338697 | 0.1377612 |
| 7 | 7316-544-T-331925 | 0.237809098 |
| 8 | 7316-2557-T-460353 | 1.228521033 |
| 9 | 7316-3208-T-555661 | 1.112207304 |
| 10 | 7316-116-T-108048 | 1.092506142 |
| 11 | 7316-126-T | 0.578837381 |
| 12 | 7316-3206-T-555638 | 1.019872041 |
| 13 | 7316-548-T-332042 | 0.750577027 |
| 14 | 7316-550-T-332072 | 0.854081823 |
| 15 | A09410-T | 0.867670293 |
| 16 | A14709-T | -0.472464998 |
| 17 | A09985-T | -0.78100549 |
| 18 | A20091-T | 0.199307053 |
| 19 | A12911-T | 1.573156211 |
| 20 | A05262-T | -1.677251826 |
| 21 | A17096-T | 1.204483478 |
| 22 | A14565-T | -1.608190321 |
| 23 | A15726-T | 0.781337902 |
| 24 | A14567-T | 1.588791241 |
| 25 | A18552-T | -0.371043758 |
| 26 | A17997-T | -1.042380139 |
| 27 | A19649-T | 0.614210096 |
| 28 | A09446-T | 1.008921009 |
| 29 | A16405-T | 1.288042308 |
| 30 | 7316-3214-T-A07083 | 0.510988205 |
| 31 | A09689-T | -0.034074539 |
| 32 | A08732-T | 0.090306155 |
| 33 | A12398-T | -0.509582327 |
| 34 | A03942-T | 0.727916426 |
| 35 | A09576-T | -0.68840405 |
| 36 | A08692-T | -1.717874371 |
| 37 | A14449-T | -0.328966631 |
| 38 | A15170-T | 1.014278335 |
| 39 | A08768-T | 0.336302279 |
| 40 | A19015-T | -0.883472479 |
| 41 | A18777-T | -0.959650971 |
| 42 | A07150-T | 1.255710685 |
| 43 | A12748-T | -1.543753607 |
| 44 | A16915-T | 0.186768419 |
| 45 | A08958-T | -1.316567616 |
| 46 | A16538-T | 0.686493025 |

B

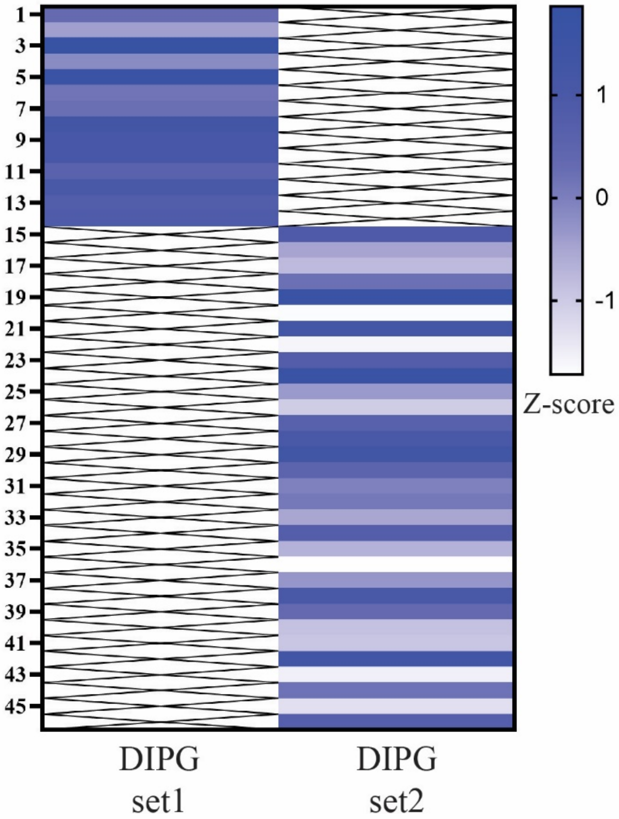

**Supplemental Figure 4. Z-score of *BIRC5* mRNA expression in DIPG.** A) Table of Z-scores of mRNA expression in patient DIPG. B) heat map of Z-scores of RNAseq data for *BIRC5* expression in human DIPG tissue, DIPG set 1 - CBTTC 1-14 (n=14), DIPG set 2- Pediatric Brain Tumor Atlas (PNOC003) 15-46 (n=31).

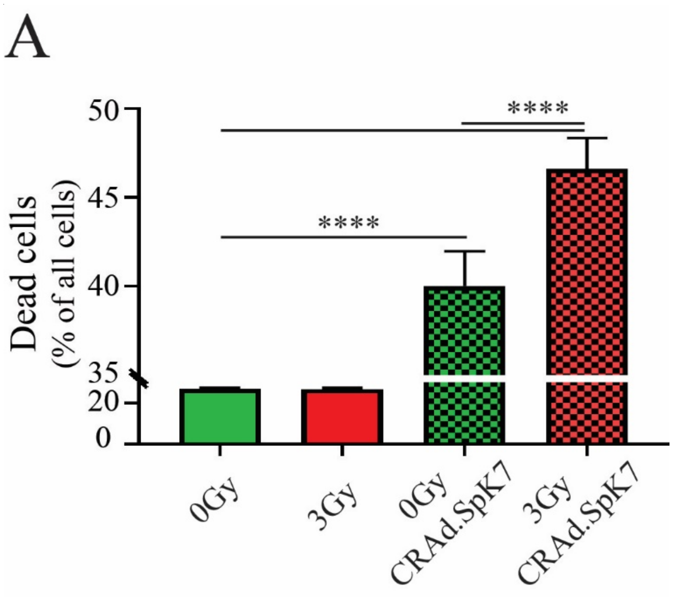


**Supplemental Figure 5. Effects of irradiation and CRAd.S.pK7 on DIPG**. A) Viability of SF8628 DIPG cells before and after 3Gy fractionated irradiation and infection with CRAd.S.K7 at MOI 10 on day 4 after infection (One-way ANOVA with Tukey's post-hoc test, n≤6. Level of significance: **** p<0.0001).

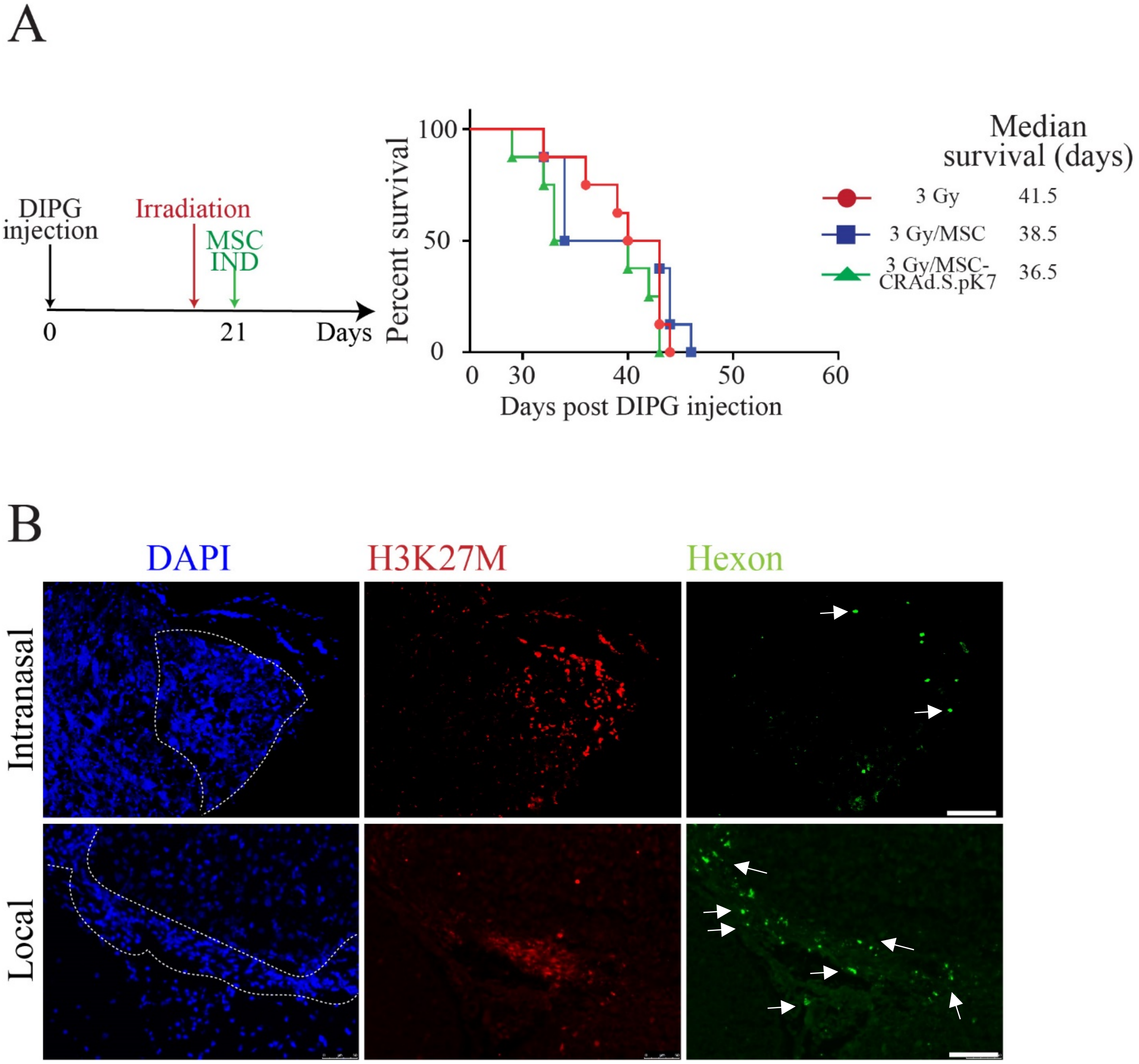


**Supplemental Figure 6. The analysis of CRAd.S.pK7 delivery by MSC to DIPG**. A) Animal treatment schema is shown on the left. Survival of DIPG bearing mice treated with IND MSC/CRAd.S.pK7 after 3Gy fractionated irradiation (Kaplan-Meier survival curves were compared using Log Rank test or Renyi test with p-values adjusted with Bonferroni correction, p=not significant (NS), n=8 per group). B) MSC-OV applied intranasally or delivered locally to DIPG in tumor-bearing mice treated with 3 Gy fractionated irradiation. DAPI- nuclear staining, H2K27M red-DIPG cells, Hexon-green-CRAd.S.pK7, indicated by white arrows. White dotted line-tumor outline. All pictures were acquired with 20X objective, white bar scale 50 µm.

**Supplemental table 1**. Z-scores for BIRC5 expression per RNAseq of human pediatric brain tumor tissue acquired from The Children's Brain Tumor Tissue Consortium (CBTTC), HGG-high grade glioma (n=132), blue table; LGG-low grade glioma (n=254), yellow table.

| Numerical order | SAMPLE_ID | Z score |
|---|---|---|
| 1 | 7316-2857-T-513841 | 0.751311736 |
| 2 | 7316-2640-T-475109 | 1.223555119 |
| 3 | 7316-1052-T-465398 | 1.263264007 |
| 4 | 7316-287-T-246895 | 1.32352493 |
| 5 | 7316-3022-T-541161 | 1.83238502 |
| 6 | 7316-3303-T-573928 | -1.055553684 |
| 7 | 7316-325-T-233387 | 0.000619035 |
| 8 | 7316-409-T-311476 | 0.423039982 |
| 9 | 7316-2092-T-407697 | 0.304686555 |
| 10 | 7316-2100-T-407801 | 1.305677338 |
| 11 | 7316-1059-T-351287 | 1.263659538 |
| 12 | 7316-2217-T-431153 | 1.154778901 |
| 13 | 7316-2554-T-472283 | 0.917730581 |
| 14 | 7316-3625-T-584791 | 1.114104497 |
| 15 | 7316-85-T-61498 | 1.382096952 |
| 16 | 7316-85-T-61659-CL-adh | -0.270257411 |
| 17 | 7316-85-T-61659-CL-susp | 1.575305475 |
| 18 | 7316-2085-T-407606 | 1.064816343 |
| 19 | 7316-515-T-331492 | 1.375905028 |
| 20 | 7316-114-T-106822 | 1.822379458 |
| 21 | 7316-466-T-323675 | 1.741883812 |
| 22 | 7316-913-T-345467 | 0.807384911 |
| 23 | 7316-913-T-345474-CL-adh | 0.015044673 |
| 24 | 7316-913-T-345474-CL-susp | 0.709901781 |
| 25 | 7316-2599-T-467328 | 0.247314617 |
| 26 | 7316-388-T-290239 | 0.360696009 |
| 27 | 7316-1936-T-386237 | 1.045484978 |
| 28 | 7316-1937-T-386252 | 1.309801537 |
| 29 | 7316-2810-T-513628 | -0.799723337 |
| 30 | 7316-3769-T-646871 | 1.049080113 |
| 31 | 7316-3521-T-576646 | 0.48710067 |
| 32 | 7316-2980-T-534980 | 0.900740102 |
| 33 | 7316-620-T-337544 | 0.424131149 |
| 34 | 7316-2610-T-467470 | 1.960393949 |

| Numerical order | SAMPLE_ID | Z score |
|---|---|---|
| 1 | 7316-2989-T-534989 | -0.375027076 |
| 2 | 7316-898-T-345214 | -1.228911679 |
| 3 | 7316-2581-T-463633 | -0.70490232 |
| 4 | 7316-3491-T-576192 | -1.123431869 |
| 5 | 7316-121-T-110375 | -1.088797108 |
| 6 | 7316-726-T-340042 | -0.021502103 |
| 7 | 7316-3504-T-576441 | -0.744217627 |
| 8 | 7316-3203-T-555602 | -0.608367786 |
| 9 | 7316-2990-T-534990 | 0.412048452 |
| 10 | 7316-2184-T-425899 | -0.651220416 |
| 11 | 7316-460-T-323574 | -0.843940677 |
| 12 | 7316-2725-T-501541 | -0.819084454 |
| 13 | 7316-1659-T-361043 | -0.619836561 |
| 14 | 7316-3627-T-584815 | -0.838909354 |
| 15 | 7316-3629-T-584839 | -0.980827775 |
| 16 | 7316-184-T-567968 | -0.974894775 |
| 17 | 7316-1745-T | -0.90140318 |
| 18 | 7316-3040-T-543985 | -0.404172693 |
| 19 | 7316-173-T-167782 | -1.212994843 |
| 20 | 7316-2154-T-409331 | -0.446781734 |
| 21 | 7316-2897-T-542852 | -0.99888631 |
| 22 | 7316-917-T-345535 | -1.04262139 |
| 23 | 7316-176-T | 0.067006295 |
| 24 | 7316-3575-T-580526 | -1.167002455 |
| 25 | 7316-1890-T-376458 | -0.880043285 |
| 26 | 7316-502-T-331285 | 1.018809589 |
| 27 | 7316-2622-T-469083 | -1.236986378 |
| 28 | 7316-2962-T-534962 | -0.980827775 |
| 29 | 7316-399-T-290428 | -1.075339088 |
| 30 | 7316-3515-T-576576 | -0.70064704 |
| 31 | 7316-1963-T-389517 | -0.627571664 |
| 32 | 7316-925-T-374499 | -0.912293469 |
| 33 | 7316-3622-T-584756 | -0.440558772 |
| 34 | 7316-1744-T-365575 | -0.869566563 |

| | | |
|---|---|---|
| 35 | 7316-1893-T | 1.197412191 |
| 36 | 7316-1893-T-376986 | 0.89211225 |
| 37 | 7316-535-T-331790 | 1.733856971 |
| 38 | 7316-1064-T-351292 | 0.792846143 |
| 39 | 7316-2152-T-409299 | 1.454882662 |
| 40 | 7316-2308-T-458799 | 1.201258751 |
| 41 | 7316-288-T-246894 | 1.578285072 |
| 42 | 7316-903-T-345298 | 1.322420661 |
| 43 | 7316-1789-T-366339 | 1.157930143 |
| 44 | 7316-445-T-317694 | 1.44350585 |
| 45 | 7316-2069-T-407398 | 0.133124493 |
| 46 | 7316-2091-T-407684 | 0.26203473 |
| 47 | 7316-2737-T-506177 | -0.311648536 |
| 48 | 7316-249-T-232228 | 1.612449849 |
| 49 | 7316-1312-T-356990 | 1.567124354 |
| 50 | 7316-851-T-344662 | 1.230169739 |
| 51 | 7316-1694-T-362487 | -0.295883297 |
| 52 | 7316-519-T-331550 | 2.076727046 |
| 53 | 7316-1187-T-356738 | 1.742549215 |
| 54 | 7316-1746-T-365613-CL-adh | 1.047027663 |
| 55 | 7316-1746-T-365613-CL-susp | 0.056871678 |
| 56 | 7316-1746-T-365608 | 1.28743638 |
| 57 | 7316-2241-T-435060 | 1.034083233 |
| 58 | 7316-1464-T-416624 | 1.738548751 |
| 59 | 7316-371-T-291950 | 2.29489722 |
| 60 | 7316-3935-T-609467 | 0.976272758 |
| 61 | 7316-1062-T-351290 | 0.275193195 |
| 62 | 7316-1723-T-364470 | 1.411057555 |
| 63 | 7316-430-T-312960 | 1.372444947 |
| 64 | 7316-716-T-339892 | 1.492014245 |
| 65 | 7316-1085-T-353519 | 0.801876404 |
| 66 | 7316-1060-T-351288 | 0.370061314 |
| 67 | 7316-1057-T-351285 | 1.483871877 |
| 68 | 7316-619-T-337543 | 0.236442211 |
| 69 | 7316-644-T-338772 | 1.168644208 |

| | | |
|---|---|---|
| 35 | 7316-2583-T-463662 | -0.639311577 |
| 36 | 7316-20-T-15915 | -0.589599408 |
| 37 | 7316-2984-T-534984 | -0.809347283 |
| 38 | 7316-375-T-242779 | -0.864377517 |
| 39 | 7316-3923-T-609299 | -1.023594115 |
| 40 | 7316-1646-T-360850 | -1.167002455 |
| 41 | 7316-70-T | -1.088797108 |
| 42 | 7316-84-T | -1.055553684 |
| 43 | 7316-884-T-344976 | -0.726513208 |
| 44 | 7316-3065-T-548496 | -0.771505263 |
| 45 | 7316-499-T-324271 | -0.651220416 |
| 46 | 7316-1866-T-373311 | -1.220914999 |
| 47 | 7316-50-T | -1.029888061 |
| 48 | 7316-3566-T-592954 | -0.986803613 |
| 49 | 7316-1760-T-365846 | -0.553293149 |
| 50 | 7316-733-T-340147 | -0.99888631 |
| 51 | 7316-264-T-232408 | -1.182051052 |
| 52 | 7316-2667-T | -0.819084454 |
| 53 | 7316-1942-T-389204 | -1.062096406 |
| 54 | 7316-2591-T-463781 | 0.35833803 |
| 55 | 7316-1851-T-373224 | -0.327719915 |
| 56 | 7316-1926-T-379786 | -0.688010917 |
| 57 | 7316-168-T-165669 | -0.61217308 |
| 58 | 7316-436-T-313084 | -1.075339088 |
| 59 | 7316-390-T-290265 | -0.515113843 |
| 60 | 7316-2689-T-485705 | -0.814201549 |
| 61 | 7316-936-T-345860 | -0.833908432 |
| 62 | 7316-3150-T-555121 | -1.197378342 |
| 63 | 7316-2729-T-501602 | -0.730903935 |
| 64 | 7316-1711-T-362742 | -1.13770205 |
| 65 | 7316-1928-T-384021 | -1.109403567 |
| 66 | 7316-3169-T-555349 | -1.13770205 |
| 67 | 7316-459-T-323557 | -0.951579148 |
| 68 | 7316-162-T-152289 | -1.182051052 |
| 69 | 7316-2174-T-425750 | -1.04262139 |

| | | |
|---|---|---|
| 70 | 7316-2151-T-409284 | -0.395320737 |
| 71 | 7316-2151-T-409293-CL-adh | 0.859254652 |
| 72 | 7316-2536-T-465316 | 1.659598442 |
| 73 | 7316-714-T-339862 | 0.657309597 |
| 74 | 7316-2594-T-463828 | 1.926834798 |
| 75 | 7316-3058-T-548405-CL-adh | 0.977949861 |
| 76 | 7316-3058-T-548405-CL-susp | 0.827051424 |
| 77 | 7316-3058-T-557515 | 1.964134362 |
| 78 | 7316-1769-T-366000 | 1.777709012 |
| 79 | 7316-1769-T-366002-CL-susp | 0.327056836 |
| 80 | 7316-392-T-290287 | -1.04262139 |
| 81 | 7316-1455-T-357608 | -0.814201549 |
| 82 | 7316-1889-T-378474 | 0.932021296 |
| 83 | 7316-440-T-313142 | 1.454568742 |
| 84 | 7316-3075-T-549171 | -0.126963775 |
| 85 | 7316-3030-T-541271 | -0.518512821 |
| 86 | 7316-2703-T-488887 | 0.810125476 |
| 87 | 7316-1774-T-366086 | -1.049062278 |
| 88 | 7316-2146-T-409210 | -0.103942948 |
| 89 | 7316-2660-T-479108 | -0.35242772 |
| 90 | 7316-2751-T-533306 | 0.575211307 |
| 91 | 7316-895-T-345161 | 1.120710632 |
| 92 | 7316-2274-T-445151 | 0.166538756 |
| 93 | 7316-1055-T-351283 | 0.153031928 |
| 94 | 7316-1099-T-353730 | 1.247285711 |
| 95 | 7316-89-T-63187 | -0.163850745 |
| 96 | 7316-1956-T-389413 | 0.917129746 |
| 97 | 7316-870-T | -0.542704731 |
| 98 | 7316-3765-T-592313 | 1.54321342 |
| 99 | 7316-2140-T-409120 | -0.43131119 |
| 100 | 7316-1946-T-389263 | 1.718022791 |
| 101 | 7316-3771-T-592398 | -0.103942948 |
| 102 | 7316-461-T-567996 | 0.686409664 |
| 103 | 7316-2561-T-460404 | -0.277862212 |
| 104 | 7316-231-T-232014 | 0.714521235 |

| | | |
|---|---|---|
| 70 | 7316-397-T-290404 | -0.659255916 |
| 71 | 7316-1681-T-361375 | -0.713479081 |
| 72 | 7316-1957-T-389427 | -1.159579535 |
| 73 | 7316-2755-T-543410 | -1.123431869 |
| 74 | 7316-1978-T-389743 | -0.722145652 |
| 75 | 7316-1658-T-361029 | -0.923328966 |
| 76 | 7316-449-T-317761 | 0.244609876 |
| 77 | 7316-1845-T-373205 | 0.169510572 |
| 78 | 7316-957-T-346215 | -1.130536205 |
| 79 | 7316-217-T | -0.963154862 |
| 80 | 7316-2744-T-507997 | -0.957346768 |
| 81 | 7316-391-T-290277 | -0.823996338 |
| 82 | 7316-896-T-345178 | -0.833908432 |
| 83 | 7316-1226-T-356777 | -1.13770205 |
| 84 | 7316-3555-T-580248 | -0.849002772 |
| 85 | 7316-93-T-64876 | -0.713479081 |
| 86 | 7316-1972-T-389653 | 1.186198419 |
| 87 | 7316-476-T-323843 | -1.04262139 |
| 88 | 7316-2291-T-447766 | -0.896011304 |
| 89 | 7316-1664-T-361118 | -0.833908432 |
| 90 | 7316-3308-T-563609 | 0.146957406 |
| 91 | 7316-612-T-337536 | -0.77614197 |
| 92 | 7316-488-T-324077 | -0.615995951 |
| 93 | 7316-1710-T-362727 | -0.906830413 |
| 94 | 7316-2669-T-479243 | -0.101881552 |
| 95 | 7316-183-T-171598 | -1.102477585 |
| 96 | 7316-1678-T-361328 | 0.596733213 |
| 97 | 7316-381-T-242851 | -0.40714451 |
| 98 | 7316-290-T-232965 | -0.912293469 |
| 99 | 7316-127-T-112912 | -1.102477585 |
| 100 | 7316-875-T-344821 | -1.011148015 |
| 101 | 7316-869-T-344719 | -0.934513595 |
| 102 | 7316-1767-T-365967 | 0.179828694 |
| 103 | 7316-97-T-66579 | -1.004994456 |
| 104 | 7316-974-T-351202 | -1.011148015 |

| | | |
|---|---|---|
| 105 | 7316-2970-T-534970 | 0.732745457 |
| 106 | 7316-204-T | 1.635477146 |
| 107 | 7316-2307-T-458786 | -1.245140631 |
| 108 | 7316-3000-T-536211 | 0.590513592 |
| 109 | 7316-624-T-337548 | 0.404266595 |
| 111 | 7316-1106-T-353836 | 1.448897726 |
| 112 | 7316-2777-T-513394 | -0.247853487 |
| 113 | 7316-2176-T-425778 | 0.866953778 |
| 114 | 7316-2176-T-425785-CL-adh | -0.917792825 |
| 115 | 7316-2756-T-508176 | 0.942579696 |
| 116 | 7316-2189-T-425973 | 0.621154585 |
| 117 | 7316-2189-T-425982-CL-adh | 1.419944715 |
| 118 | 7316-1844-T-373190 | -0.462546973 |
| 119 | 7316-343-T-242379 | 0.38391352 |
| 120 | 7316-238-T-232096 | 1.458327946 |
| 121 | 7316-255-T-232300 | 1.416006643 |
| 122 | 7316-622-T-337546 | 0.901965342 |
| 123 | 7316-195-T-202630 | 1.142052566 |
| 124 | 7316-195-T-202952-CL-adh | 1.300398459 |
| 125 | 7316-195-T-203029-CL-susp | 2.278700929 |
| 126 | 7316-3122-T-550919 | 1.313905287 |
| 127 | 7316-2288-T-465298 | 1.081245424 |
| 128 | 7316-1656-T-360998 | 0.858609814 |
| 129 | 7316-161-T | 1.312415367 |
| 130 | 7316-1763-T-365898 | 2.267019013 |
| 131 | 7316-1763-T-365902-CL-adh | 1.398721758 |
| 132 | 7316-1763-T-365902-CL-susp | -0.833908432 |
| 133 | 7316-1068-T-351296 | 1.179659143 |

| | | |
|---|---|---|
| 105 | 7316-891-T-345093 | -1.068691261 |
| 106 | 7316-329-T-234369 | -1.159579535 |
| 107 | 7316-2215-T-431127 | -0.859220803 |
| 108 | 7316-882-T-344942 | -1.159579535 |
| 109 | 7316-482-T-323958 | NA |
| 110 | 7316-175-T-168637 | 0.524637771 |
| 111 | 7316-3055-T-548356 | -0.969004003 |
| 112 | 7316-1635-T-360684 | -1.159579535 |
| 113 | 7316-2195-T-426065 | -1.023594115 |
| 114 | 7316-924-T-345654 | -0.963154862 |
| 115 | 7316-134-T-115887 | -0.106009491 |
| 116 | 7316-2664-T-479169 | -0.794953007 |
| 117 | 7316-35-T-27190 | NA |
| 118 | 7316-1081-T-353459 | -0.288111879 |
| 119 | 7316-1087-T-353550 | -1.109403567 |
| 120 | 7316-32-T-25911 | -0.992822913 |
| 121 | 7316-1975-T-389698 | -1.167002455 |
| 122 | 7316-3922-T-609284 | -0.744217627 |
| 123 | 7316-1134-T-356677 | 0.825710013 |
| 124 | 7316-3140-T-551166 | -0.874788346 |
| 125 | 7316-77-T | -1.167002455 |
| 126 | 7316-470-T-323744 | -0.226040072 |
| 127 | 7316-3154-T-555169 | -0.532250066 |
| 128 | 7316-1661-T-361075 | -1.109403567 |
| 129 | 7316-1660-T-361058 | -0.906830413 |
| 130 | 7316-2285-T-465290 | -0.635379741 |
| 131 | 7316-932-T-345792 | -0.945851444 |
| 132 | 7316-332-T-242248 | -1.130536205 |
| 133 | 7316-207-T-212603 | -0.70917959 |
| 134 | 7316-1086-T-353536 | -0.643262181 |
| 135 | 7316-3919-T-609243 | -1.082040744 |
| 136 | 7316-741-T-341931 | -0.940163105 |
| 137 | 7316-265-T-232422 | -1.062096406 |
| 138 | 7316-487-T-324062 | -1.20514976 |
| 139 | 7316-152-T-148056 | -0.753213911 |

| | | |
|---|---|---|
| 140 | 7316-2752-T-508116 | NA |
| 141 | 7316-1679-T-361343 | -1.109403567 |
| 142 | 7316-1652-T-360938 | -0.896011304 |
| 143 | 7316-3074-T-549156 | 0.423039982 |
| 144 | 7316-2324-T-459010 | -1.197378342 |
| 145 | 7316-3554-T-580234 | -0.150643284 |
| 146 | 7316-3062-T-548455 | -0.36083033 |
| 147 | 7316-1088-T-353564 | -0.238087592 |
| 148 | 7316-2589-T-463752 | -0.179530645 |
| 149 | 7316-1648-T-360878 | -0.77614197 |
| 150 | 7316-1953-T-389367 | -0.095727992 |
| 151 | 7316-222-T-225384 | -1.189679216 |
| 152 | 7316-874-T-344804 | -1.055553684 |
| 153 | 7316-437-T-313099 | -0.945851444 |
| 154 | 7316-393-T-290301 | -0.819084454 |
| 155 | 7316-1084-T-353504 | -0.036584981 |
| 156 | 7316-666-T-339122 | -1.055553684 |
| 157 | 7316-1113-T-353939 | -1.004994456 |
| 158 | 7316-2249-T-435163 | -0.683841387 |
| 159 | 7316-186-T-172877 | -0.184065725 |
| 160 | 7316-350-T-242463 | -1.109403567 |
| 161 | 7316-944-T-345996 | -0.880043285 |
| 162 | 7316-1803-T-366682 | -1.130536205 |
| 163 | 7316-206-T-212179 | -0.401211509 |
| 164 | 7316-475-T-323828 | -1.20514976 |
| 165 | 7316-275-T-232540 | -0.462546973 |
| 166 | 7316-880-T-344906 | NA |
| 167 | 7316-370-T-242720 | -1.029888061 |
| 168 | 7316-2172-T-456310 | -0.945851444 |
| 169 | 7316-2075-T-407476 | -1.01734767 |
| 170 | 7316-267-T-232444 | -0.567625109 |
| 171 | 7316-3061-T-548439 | -0.597055765 |

| | | |
|---|---|---|
| 172 | 7316-177-T | NA |
| 173 | 7316-2197-T-430377 | -1.20514976 |
| 174 | 7316-1958-T-389443 | -0.945851444 |
| 175 | 7316-2135-T-408257 | -0.748703545 |
| 176 | 7316-1109-T-353881 | -0.934513595 |
| 177 | 7316-1943-T-389219 | -1.220914999 |
| 178 | 7316-3076-T-549180 | -1.011148015 |
| 179 | 7316-923-T-345637 | -0.828937547 |
| 180 | 7316-692-T-339532 | -0.969004003 |
| 181 | 7316-2566-T-462544 | -0.639311577 |
| 182 | 7316-368-T-242681 | -0.854096019 |
| 183 | 7316-462-T-323607 | 0.018612135 |
| 184 | 7316-2310-T-458825 | -1.182051052 |
| 185 | 7316-946-T-346029 | -0.309000053 |
| 186 | 7316-1668-T-361179 | -0.99888631 |
| 187 | 7316-154-T-148904 | 0.015044673 |
| 188 | 7316-3570-T-580457 | -0.040399043 |
| 189 | 7316-562-T-336223 | -0.730903935 |
| 190 | 7316-663-T-339077 | -1.049062278 |
| 191 | 7316-1977-T-389729 | -0.627571664 |
| 192 | 7316-297-T-233051 | -0.546219174 |
| 193 | 7316-2642-T-475135 | NA |
| 194 | 7316-954-T-346166 | -0.828937547 |
| 195 | 7316-9-T-7029 | -0.804521322 |
| 196 | 7316-2275-T-445164 | -0.951579148 |
| 197 | 7316-136-T-552494 | -0.874788346 |
| 198 | 7316-262-T-258405 | 0.085270125 |
| 199 | 7316-2632-T-469213 | -0.790210014 |
| 200 | 7316-1805-T-366712 | -0.3496457 |
| 201 | 7316-362-T-242608 | -0.864377517 |
| 202 | 7316-643-T-338757 | -0.917792825 |
| 203 | 7316-344-T-242392 | -0.880043285 |

| | | |
|---|---|---|
| 204 | 7316-442-T-315526 | -0.980827775 |
| 205 | 7316-1794-T-366549 | -0.539205149 |
| 206 | 7316-2899-T-542859 | -1.182051052 |
| 207 | 7316-677-T-339287 | -0.61217308 |
| 208 | 7316-516-T-331505 | -0.659255916 |
| 209 | 7316-247-T-232204 | -0.957346768 |

| | | |
|---|---|---|
| 210 | 7316-146-T-145834 | 0.055170444 |
| 211 | 7316-119-T | -1.109403567 |
| 212 | 7316-361-T-242595 | -1.13770205 |
| 213 | 7316-681-T-339367 | -0.422165793 |
| 214 | 7316-33-T | NA |
| 215 | 7316-2663-T-479154 | -0.647231733 |
| 216 | 7316-235-T-232061 | -0.70490232 |
| 217 | 7316-3054-T-548341 | -0.560428107 |
| 218 | 7316-3068-T-548538 | -1.068691261 |
| 219 | 7316-2130-T-408191 | -0.986803613 |
| 220 | 7316-400-T-290442 | -0.874788346 |
| 221 | 7316-2167-T-425644 | -0.316971062 |
| 222 | 7316-193-T | -1.182051052 |
| 223 | 7316-472-T-323777 | -1.011148015 |
| 224 | 7316-642-T-338742 | -1.049062278 |
| 225 | 7316-1949-T-389307 | -0.880043285 |
| 226 | 7316-1095-T-353669 | -0.207121534 |
| 227 | 7316-1643-T-360803 | -0.880043285 |
| 228 | 7316-2099-T-407788 | -0.179530645 |
| 229 | 7316-167-T | -1.004994456 |
| 230 | 7316-3522-T-576658 | -0.739755891 |
| 231 | 7316-148-T-146367 | -1.004994456 |
| 232 | 7316-1090-T-353596 | -0.980827775 |
| 233 | 7316-345-T-242405 | -0.504999868 |
| 234 | 7316-1779-T-366169 | -1.004994456 |
| 235 | 7316-3762-T-592273 | -0.389471596 |
| 236 | 7316-3751-T-592117 | -1.174492552 |
| 237 | 7316-2006-T-391850 | -0.874788346 |
| 238 | 7316-495-T-324198 | -1.095609081 |
| 239 | 7316-3556-T-580262 | -1.220914999 |
| 240 | 7316-2572-T-463497 | -0.532250066 |
| 241 | 7316-2147-T-409224 | -0.571247215 |

| | | |
|---|---|---|
| 242 | 7316-2193-T-426035 | 0.245963351 |
| 243 | 7316-485-T-347672 | -1.212994843 |
| 244 | 7316-196-T-203061 | -0.744217627 |

| | | |
|---|---|---|
| 245 | 7316-2204-T-430468 | -0.890654328 |
| 246 | 7316-2208-T-430520 | -0.928902387 |
| 247 | 7316-938-T-345894 | -0.525352943 |
| 248 | 7316-1683-T-361403 | -1.152222598 |
| 249 | 7316-441-T-314920 | -0.491704 |
| 250 | 7316-658-T-338982 | -0.157220669 |
| 251 | 7316-554-T-332455 | -1.13770205 |
| 252 | 7316-2305-T-458760 | -1.123431869 |
| 253 | 7316-934-T-345825 | -0.52879432 |
| 254 | 7316-2245-T-435112 | -0.696413527 |
| 255 | 7316-3323-T-564960 | -0.137643278 |
| 256 | 7316-2753-T-508131 | -0.726513208 |
| 257 | 7316-2988-T-534988 | -1.189679216 |
| 258 | 7316-346-T-242416 | -0.986803613 |
| 259 | 7316-3139-T-551154 | -1.109403567 |
| 260 | 7316-117-T-108479 | -0.688010917 |
| 261 | 7316-2723-T-501511 | -1.049062278 |
